# Notch1 Phase Separation Coupled Percolation facilitates target gene expression and enhancer looping

**DOI:** 10.1101/2023.03.17.533124

**Authors:** Gregory Foran, Ryan Douglas Hallam, Marvel Megaly, Anel Turgambayeva, Daniel Antfolk, Yifeng Li, Vincent C. Luca, Aleksandar Necakov

## Abstract

The Notch receptor is a pleiotropic signaling protein that translates intercellular ligand interactions into changes in gene expression *via* the nuclear localization of the Notch intracellular Domain (NICD). Using a combination of immunohistochemistry, RNA *in situ,* Optogenetics and super-resolution live imaging of transcription in human cells, we show that the N1ICD can form condensates that positively facilitate Notch target gene expression. We determined that N1ICD undergoes Phase Separation Coupled Percolation (PSCP) into transcriptional condensates, which recruit, enrich, and encapsulate a broad set of core transcriptional proteins. We show that the capacity for condensation is due to the intrinsically disordered transcriptional activation domain of the N1ICD. In addition, the formation of such transcriptional condensates acts to promote Notch-mediated super enhancer-looping and concomitant activation of the MYC protooncogene expression. Overall, we introduce a novel mechanism of Notch1 activity in which discrete changes in nuclear N1ICD abundance are translated into the assembly of transcriptional condensates that facilitate gene expression by enriching essential transcriptional machineries at target genomic loci.

## Introduction

Notch receptors constitute a family of signalling proteins that translate ligand-mediated activation by neighbouring cells at sites of direct, intercellular contact to changes in gene expression.^1–3^ The Notch signalling pathway can be viewed as an integrative molecular counter of productive cellular interactions, which translates these interactions into changes in cell-type specific target gene expression.^1–5^ Notch signals are used iteratively at a wide range of distinct, context-dependent cellular decision points, and drive transcriptional programs that are highly sensitive to gene dosage.^1,6^ Both deficiencies and slight perturbations of Notch signalling are associated with developmental abnormalities and numerous diseases, including T-Cell Acute Lymphoblastic Leukemia(T-ALL) in humans.^2–5,7–14^

The intracellular domain of the Notch receptor (NICD) is the primary effector of Notch signalling and is released from the membrane through proteolytic cleavage by the gamma secretase complex in response to ligand-based activation.^3,15^ Cleavage liberates the NICD from the plasma membrane, resulting in nuclear translocation.^1–3,15^ Nuclear NICD physically associates with its DNA binding partner RBPJ at discrete genomic RBPJ-binding sites.^5,16–18^ In concert with additional factors, including MAML1, p300, and other core transcriptional machinery, NICD drives assembly of the Notch transcriptional activation complex, thereby activating Notch target gene expression.^3,6,19,20^ A fundamental unresolved question regarding Notch signalling is how increases in nuclear NICD abundances regulate enhancer looping, and how they are translated into discrete changes in transcriptional output across multiple target genes.

Recent investigations have demonstrated that enhancer-looping is regulated by a variety of proteins that undergo a process, which has been recently defined to properly account for the variety of features specific to protein phase separation, termed Phase Separation Coupled Percolation (PSCP), to assemble into membraneless organelles termed biomolecular condensates.^21–24^ Biomolecular condensates are dynamic, motile, self-organizing structures that can spontaneously form, exhibit varying degrees of mixing based on their fluidity, and have the capacity to undergo homotypic fusion.^25–27^ The ability of proteins to undergo PSCP into biomolecular condensates is driven by intrinsically disordered regions(IDRs) that they possess.^28,29^ The formation of a specialized subset of nuclear biomolecular condensates, termed transcriptional condensates, has previously been linked to the regulation of target gene expression, enhancer looping, and increased local concentrations of transcriptionally active proteins.^30,31^ In addition, several transcriptional regulators, including YAP, TAZ, MED1, P300, and BRD4 have been shown to phase separate into transcriptional condensates.^30–34^

Previous studies have provided strong evidence that the C-terminal domain of Notch1 is essential to drive high levels of Notch target gene expression in multiple contexts.^35,36^ However, the mechanisms through which the C-terminal tail of Notch1 potentiates gene expression has not been resolved. Here we performed *in silico* analysis of the Notch1 NICD(N1ICD) and identified an IDR in the C-terminal tail using multiple predictive models, which we tested using molecular dynamics simulation (MDS).^37,38^ Using purified N1ICD we demonstrate that the N1ICD undergoes PSCP in a salt- and concentration-dependent manner.

To study the capacity of N1ICD to form functional transcriptional condensates *in vivo* we began by generating several novel molecular tools to simultaneously control and monitor the activity of Notch1 in living human cells. The first of these tools is an engineered Optogenetic Notch protein construct(OptoNotch) that provides the ability to precisely titrate intranuclear levels of transcriptionally active N1ICD in real-time. By employing Opto-Notch, we show for the first time that the N1ICD spontaneously self-organizes into dynamic, transcriptionally active condensates with liquid-like properties, which recruit and enrich several key factors necessary for transcriptional activation of canonical Notch1 target genes. We also showed that Notch1 transcriptional condensates exhibit dynamic growth and shrinkage, that spontaneous Notch1 condensate self-assembly is dependent upon the presence of the intrinsically disordered C-terminal N1ICD tail, and that the Notch1 ankyrin repeats are responsible for seeding Notch1 condensate formation at target gene loci. We then demonstrated that our OptoNotch tool can function within the range of endogenous Notch signalling both in HEK293 and in T-ALL cells (CUTTL1), highlighting that OptoNotch is a functional model system that allows for the study of Notch dynamics in living cells.

We further investigated the relationship between signal activation and transcription using a novel fluorescent Notch transcriptional reporter system that we developed, which provides a high-fidelity, quantitative, temporal read-out of the formation of nascent transcriptional foci of the Notch1 target gene Hes1 in live cells. By employing this tool, we uncovered a novel regulatory mechanism in which N1ICD self-associates into intranuclear liquid spherical shell condensates that resist transcription-driven dissociation, exhibit dynamic changes in volume and content, and which thereby increase the duration of transcriptional bursting in a concentration-dependent manner.

In addition, we demonstrate that Notch1 transcriptional condensate assembly promotes super-enhancer looping between the Notch-Dependant *MYC* Enhancer (NDME) and the *MYC* promoter, located >1.7 megabases away, and facilitates concomitant expression of the *MYC* protooncogene. This establishes a novel mechanism of Notch1-mediated super-enhancer looping in human T-ALL cells *via* condensate formation and provides valuable insights into the mechanisms by which Notch signalling regulates target gene expression.^7–9,39^

## Results

### The human N1ICD exhibits properties consistent with PSCP

To assess the potential of the N1ICD to undergo PSCP, we first performed *in silico* analysis on the human N1ICD to predict disordered regions and sequence features known to drive PSCP.^25,37,38,40–43^ Consistent with a previous study that identified an IDR in the Notch1 RBPJ-associated motif(RAM) for transcriptional activation complex assembly through charge-patterning-mediated Notch1/RBPJ interaction,^44^ analyses using available protein disordered prediction tools (IUPRED, ALPHAFOLD, GROMACS, etc.) also provided preliminary evidence showing that the carboxy-terminal tail containing the transcriptional activation domain(TAD), which plays a critical role in Notch1-mediated transcriptional activation, contains sequences that exhibit features consistent with a high degree of structural disorder, which is common to classical IDR’s(Figure 1A/B, Extended Data Figure 1).^44,45^ Next, we used MDS analysis to calculate the root mean square deviation (RMSD) values over time per residue, and observed that, consistent with our disorder prediction results, the N1ICD TAD domain (AA2120-2555) has a significantly higher RMSD value and corresponding degree of motility and disorder, than either the Full-length N1ICD (AA1754-2555) or the Ram-Ankyrin domain alone (AA1754-2119) (Extended data figure 1A,B), providing further evidence that the N1ICD TAD domain is disordered. Collectively, these results led us to hypothesize that N1ICD could potentially undergo PSCP to form condensates.

**Figure 1:**
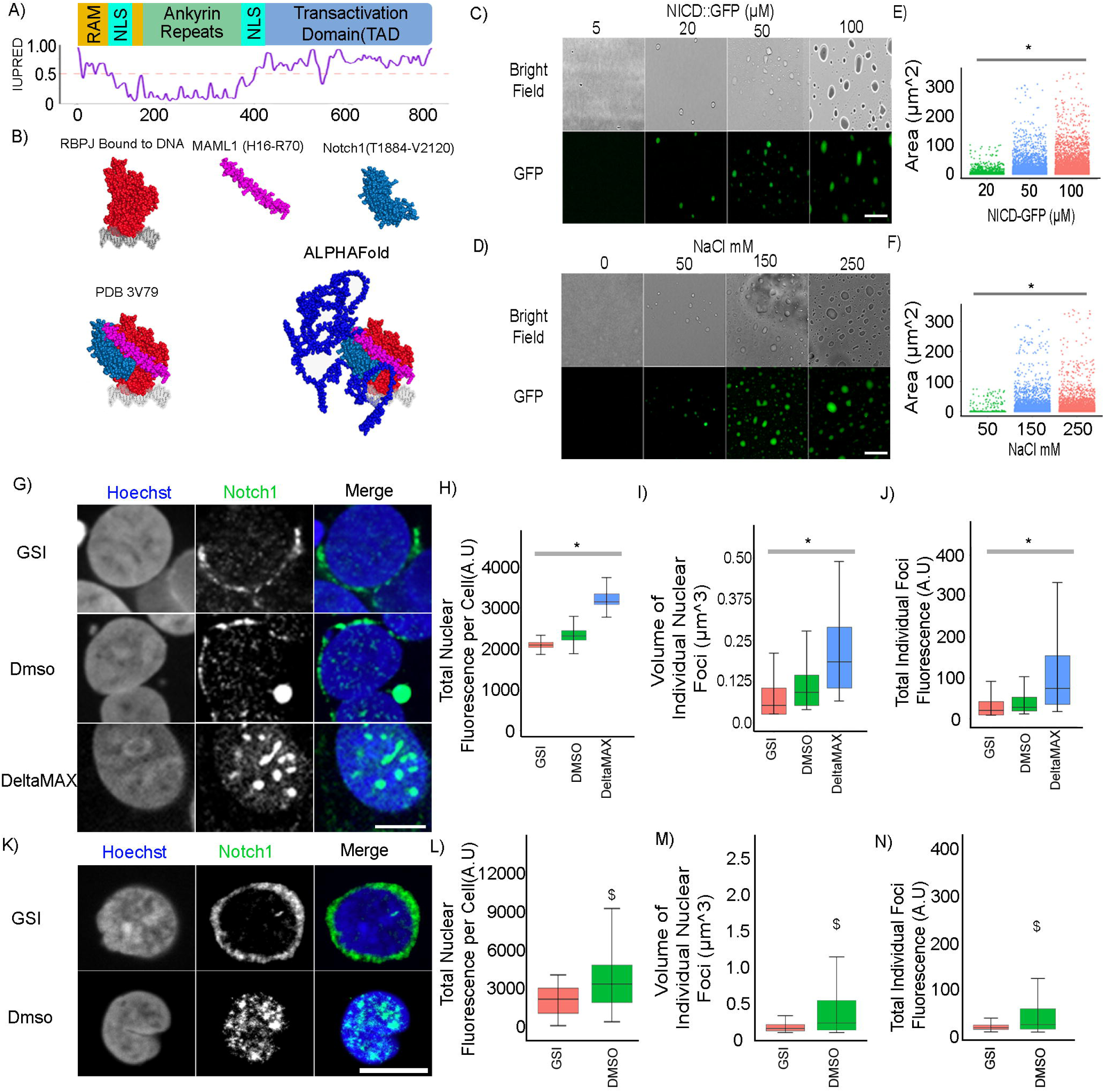
The Human Notch1 Intracellular Domain Exhibits Condensate/PSCP Behavior Both In Isolation And In Cells. A) N1ICD sequence analysis using prediction of intrinsic disorder by IUPRED. B) Individual structures of RBPJ (red), MAML1(purple) and Notch1(teal) (all taken from PDB 3V79) which form the Notch 1 activation complex, whose structure is represented by PDB 3V79(bottom left), onto which the AlphaFold structure prediction of the N1ICD TAD (dark blue) is super-imposed (bottom right). C) Droplet assay with increasing concentration of purified N1ICD::GFP protein at 150mM NaCl. Brightfield(top), green fluorescence(bottom). D) Droplet assay with increasing NaCl concentration with 50μM NICD::GFP protein. Brightfield(top), green fluorescence(bottom). E) Area of condensates with N1ICD::GFP concentration of 20μM, 50μM and 100μM forming droplets with an average area of 1.69μm^2^(+/-6.09), 9.35μm^2^(+/-21.1), and 16.1μm^2^(+/-31.1) respectively. N=5000 foci per condition. F) Area of condensates with NaCl concentration of 50 mM, 150 mM and 250mM forming droplets with an average area of 5.34μm^2^(+/-14.6), 8.49μm^2^(+/-22.9), and 12.4μm^2^(+/-27.9). N=500 foci per condition. G) Endogenous Notch1 fluorescence immunostainings in HEK293 cells treated with GSI (top), DMSO (middle), or plated on DeltaMAX(bottom). H) Total amount of Nuclear Notch1 protein per cell from panel G showing GSI, DMSO, and DeltaMAX treated cells have 2107(+/-124), 2367(+/-209), and 3260(+/-319) A.U of Notch1, respectively. N= 4000 cells per condition. I) Volume of individual nuclear foci from panel M showing GSI, DMSO, and DeltaMAX treated cells form Notch1 foci that are on average 0.092μm^3^(+/-0.234), 0.145μm^3^(+/-0.229), and 0.326μm^3^ (+/-0.370), respectively. N= 47000 foci per condition. J) Total Fluorescence per individual nuclear foci from panel M showing GSI, DMSO, and DeltaMAX treated cells have foci of 55(+/-126), 67 (+/-151), and 192(+/-738) A.U of Notch1. N= 47000 foci per condition. K) Endogenous Notch1 fluorescence immunostainings in T-ALL cells treated either with DMSO or GSI. L) Total amount of Nuclear Notch1 protein per cell from panel M showing DMSO and GSI treated cells have 3648(+/-2607) and 1989(+/-1107) A.U of Notch1, respectively. N= 12000 cells per condition. M) Volume of individual nuclear foci from panel M showing DMSO and GSI treated cells form Notch1 foci that are on average 0.672μm^3^(+/-0.532) and 0.174μm^3^(+/-0.0585) respectively. N= 16000 foci per condition. N) Total Fluorescence per individual nuclear foci from panel M showing DMSO and GSI treated cells have foci of 49.4(+/-55.3) and 19 (+/-6.7) A.U of Notch1. N= 16000 foci per condition. C/D scale bar = 50μm. G/K scale bar = 10 μm. *p<0.01 One-way ANOVA+Tukey post-hoc. $ p<0.01 on a student t-test. See Source Data Figure 1

As an initial test of this hypothesis, we titrated and imaged isolated N1ICD::GFP protein in solution, which demonstrated the formation of phase-separated droplets starting at a concentration of 20μM, where with increasing concentration, we observed a concomitant increase in the average size of N1ICD::GFP foci(Figure 1C/E). As evidence that polar/ionic interactions play an important role in NICD::GFP PCPS, N1ICD phase separation exhibited a dependence upon salt concentration, where condensate size was proportional to the amount of salt present, while we observed no N1ICD condensate formation in the absence of salt(Figure 1D/F). Consistent with previous studies that reported the formation of hollow condensates at elevated salt concentrations, intra-condensate cavities could be observed at higher NaCl concentrations(Extended Data Figure 2A).^46,47^ We also observed a significant decrease in N1ICD::GFP droplets after treatment with 1,6-hexanediol, a well-characterized aliphatic alcohol commonly used to disrupt biomolecular condensates (Extended Figure 2B).^48^

**Figure 2:**
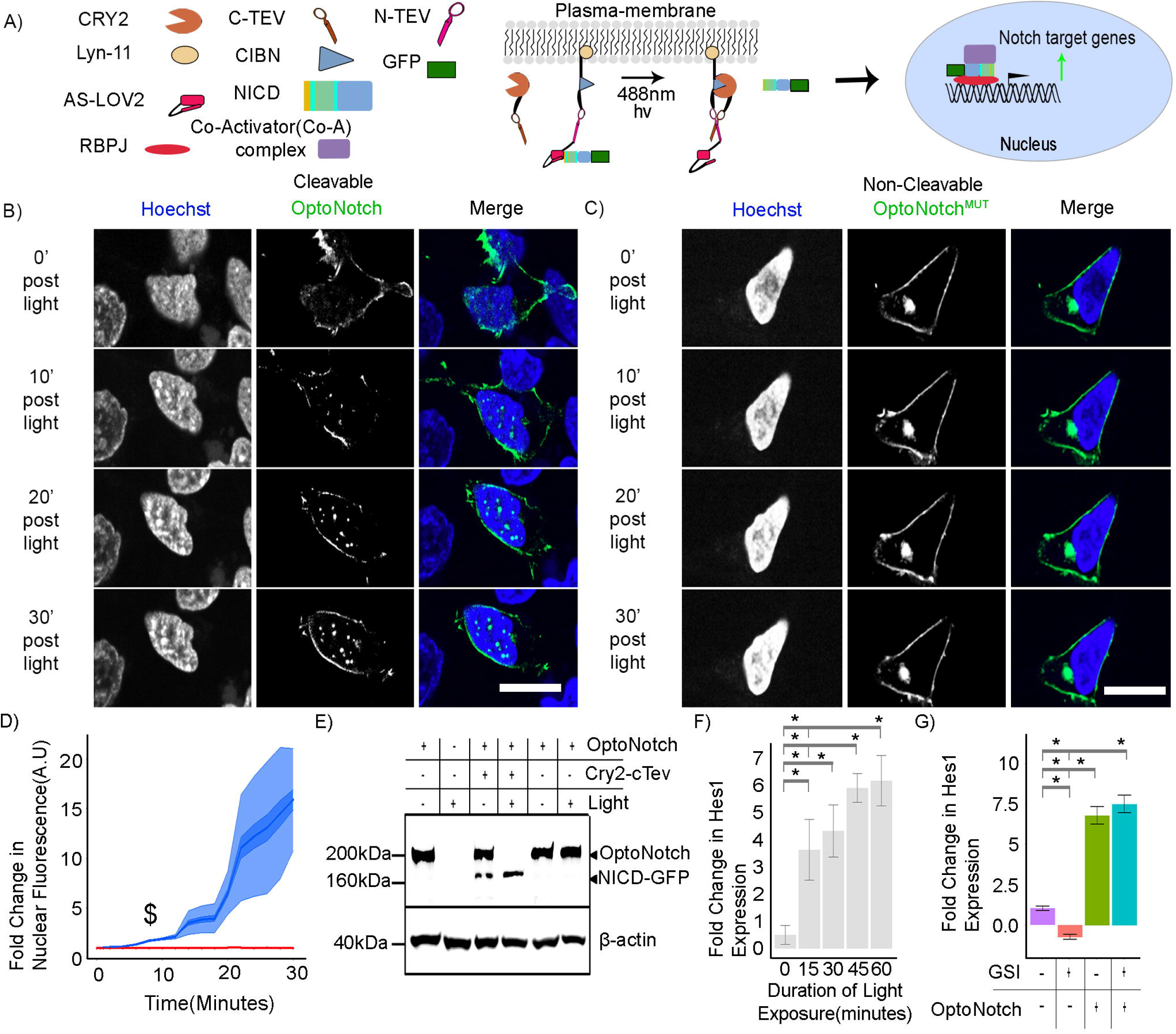
Development Of An Optogenetic Tool To Control Notch1. A) Schematic of Notch1 nuclear localization following light activation with the OptoNotch expression construct. B) Time series of OptoNotch activation in HEK293 cells. C) Time series of OptoNotch^mut^ activation in HEK293 cells. D) OptoNotch (Blue) nuclear localization compared to OptoNotch^mut^ (red) central line-mean, dark blue-standard error of the mean (SEM), and the light blue-standard deviation (SD). N = 50 cells per condition. $ p<0.01 on a student t-test. E) Western blot of OptoNotch activation in HEK293 cells under various conditions. F) qPCR of Hes1 with varying durations of OptoNotch activation with fold changes in Hes1 expression of 1.525(+/-1.342) 3.690(+/-1.134) 4.390(+/-0.972) 6.00(+/-0.535) 6.27(+/-0.941) for 0,15, 30, 45, and 60 minutes of light activation, respectively. G) qPCR of HES1 with or without GSI treatment and with or without OptoNotch activation showing a relative fold expression of Hes1 of 1.00 (+/- 0.109), 6.78(+/- 0.540), -0.737(+/- 0.084), and 7.491(+/- 0.549) for wildtype + vehicle, OptoNotch + vehicle, Wildtype + GSI, and OptoNotch + GSI respectively. C/D scale bar 10μm. *p<0.01 One-way ANOVA+Tukey post-hoc. All data acquired in live HEK293 cells. See Source Data Figure 2

### Endogenous N1ICD has the capacity to form intranuclear condensates

To address whether N1ICD forms condensates within the working range of endogenous Notch1 levels, we then investigated the effects of inhibiting endogenous Notch signalling, through the application of a gamma secretase inhibitor (GSI; compound E), or ectopically activating Notch signalling, through the addition of surface-immobilized Notch1 ligand (DeltaMAX), on the total abundance of Notch1 within the nucleus as well as the relative size and intensity of Nuclear Notch1 condensates (Figure1G, Extended Data Figure3:A/E).^49^ Following the stimulation of endogenous Notch1 with DeltaMAX we observed a significant increase in total Nuclear Notch1 that correlated with a significant increase in large intranuclear N1ICD foci, which we did not observe with either GSI or DMSO treatment (Figure1H-J, Extended Data Figure3:B-D,F-H). We then extended our analysis to include a human T-ALL cancer cell line (CUTTL1) that is sensitive to GSI treatment, and in which Notch1 is both constitutively cleaved and activated in a ligand-independent manner, and in which both endogenous Notch1 activity and nuclear abundance are high(Figure1:K, Extended Data Figure 3:I/M)^7–9,50–52^ . Using fluorescence immunohistochemistry, we observed a significant nuclear abundance of N1ICD that was organized into large intranuclear foci, which disappeared upon GSI inhibition(Figure1:L-N, Extended Data Figure 3:J-L/N-P). To validate that both our GSI and DeltaMAX treatments were effective in either inhibiting endogenous Notch1 activity or increasing Notch1 activity, we performed Western blotting for cleaved, activated Notch1. As anticipated, we observed that HEK293 cells exhibit a basal level of endogenous Notch1 activity, which is inhibited by GSI treatment, and which increases in response to DeltaMAX ligand activation(Extended Data Figure 4), demonstrating the functionality and utility of this treatment strategy.

### Development of OptoNotch: an Optogenetic Tool to Control Notch activity

To further characterize N1ICD condensates in living human cells, we next developed a novel tool that affords us the ability to perform pulse-chase experiments on N1ICD nuclear translocation under light-gated control. To do so, we adapted an existing Split Tobacco Etch Virus (TEV) protease-based optogenetic cleavage system to generate transgenic constructs that provide precise light-gated control over the release of ectopically expressed N1ICD from the plasma membrane upon light exposure; henceforth referred to as OptoNotch(Figure 2A).^53–57^ Cells expressing OptoNotch demonstrate a significant, titratable, increase in nuclear N1ICD signal, predominantly localized to discrete nuclear foci in response to illumination with blue light (∼490 nm)(Figure 2B, Movie 1). As a control for non-specific cleavage and aberrant nuclear localization, we designed an OptoNotch construct with a point mutation in a key residue in the canonical TEV cleavage sequence essential for TEV-mediated cleavage, named OptoNotch^mut^.^55^ In contrast to OptoNotch, OptoNotch^mut^ exhibits no increase in Nuclear N1ICD, remaining tethered to the plasma membrane despite continuous blue light illumination(Figure 2C). We next measured the kinetics of light-induced N1ICD nuclear translocation with OptoNotch and observed a significant increase in N1ICD within the nucleus over 30 minutes with a concomitant appearance of prominent nuclear N1ICD foci starting at 8 minutes following activation with blue light(Figure 2D). In contrast, OptoNotch^mut^ does not exhibit an observable accumulation of N1ICD in the nucleus regardless of illumination status or duration(Figure 2D), demonstrating that Opto-Notch provides precise, titratable, light-gated control over the nuclear translocation of N1ICD. OptoNotch cleavage was then validated using Western blotting, where we observed a light-gated cleavage of full-length, plasma membrane-tethered OptoNotch, resulting in the production of the expected ∼160 kDa fragment following exposure to blue light(Figure 2E). Following blue light activation of OptoNotch, we observed a concomitant increase in Hes1 expression in HEK293 cells over the course of one hour(Figure 2F). To test the orthogonality of OptoNotch with respect to endogenous Notch signalling, we pharmacologically inhibited endogenous Notch activity with a gamma-secretase inhibitor(GSI) with simultaneous OptoNotch activation. Consistent with OptoNotch function being orthogonal to endogenous Notch activity, OptoNotch is insensitive to GSI treatment and can rescue expression of Hes1; a direct Notch target gene, independent of endogenous Notch activity(Figure 2G).^1,2,5,15^ We then sought to compare the formation of Notch1 condensates at endogenous levels between untransformed HEK293 cells activated with DeltaMAX, uninhibited T-ALL cells, and HEK293 cells following OptoNotch activation(Extended Data Figure 5A). First, we demonstrated that both OptoNotch and DeltaMAX can drive Notch1 target gene expression (Hes1) significantly over baseline, where we observed that the addition of exogenous ligand drives Hes1 expression above that of OptoNotch (Extended Data Figure 5B). Following OptoNotch activation, we observed a significant increase in nuclear N1ICD abundance compared to endogenous N1ICD in both DeltaMAX-treated HEK293 cells and T-ALL cells (Extended Data Figure 5C). We also observed that T-ALL cells form the largest N1ICD condensates, which are significantly larger than condensates we observed in both DeltaMAX-treated and OptoNotch-activated HEK293 cells(Extended Data Figure 5D). However, we also observed that OptoNotch forms condensates with a higher signal intensity, on average, compared to T-ALL and DeltaMAX-treated HEK293 cells(Extended Data Figure 5E). In addition, we also observed that T-ALL cells form significantly fewer condensates compared to OptoNotch and DeltaMAX, with OptoNotch activation resulting in the highest number amongst these conditions (Extended Data Figure 5F). With regard to the total range of volumes and number of condensates per cell we observe that OptoNotch works within the range of endogenous Notch1 condensates formed from DeltaMAX treated and untreated T-ALL cells, compared to total nuclear intensity and individual intensity were there are several data points outside of the working range of both DeltaMAX treated or untreated T-ALL cells. Taken together, these data demonstrate that OptoNotch condensates show similar characteristics to those formed by endogenous Notch1 at high levels of Notch activity, thus providing evidence that OptoNotch can serve as a useful model tool to study N1ICD condensate dynamics.

Overall, our data establishes OptoNotch as a functional light-gated tool capable of regulating Notch activity through the nuclear localization of N1ICD, and concomitant expression of the Notch target gene Hes1, even in the absence of endogenous Notch signalling.

### Biophysical characterization of N1ICD nuclear foci

To characterize the biophysical properties of nuclear N1ICD condensates, we initially used Fluorescence Recovery After Photo-bleaching(FRAP).^27,58–60^ Following complete photo-bleaching, we observed that individual intranuclear N1ICD foci exhibited a dynamic recovery of nearly half of the total signal over the course of 5 minutes (Figure 3A/B, Movie 2), demonstrating that N1ICD condensates have liquid-like properties, and providing further evidence for PSCP. In addition, we observed frequent instances of intra-nuclear N1ICD condensate fusion, further demonstrating the liquid-like properties of N1ICD condensates, and providing strong evidence for PSCP(Figure 3C, Movie 3). ^21,26,27^ Interestingly, by quantifying the intensity of individual condensates, we observed that the fluorescence of post-fusion condensates equates to the sum of the fluorescence of the individual pre-fusion condensates(Figure 3J), demonstrating that N1ICD levels are maintained during condensate fusion. To further test whether nuclear N1ICD foci undergo PSCP, we first treated OptoNotch-activated HEK293 cells with 5% 1,6-Hexanediol; a disruptor of biomolecular condensation, and observed a near-complete loss of all nuclear foci immediately (∼6 seconds) following treatment(Extended Data Figure 2C-E), providing further evidence supporting the hypothesis that intranuclear N1ICD foci undergo PSCP to form biomolecular condensates.

**Figure 3:**
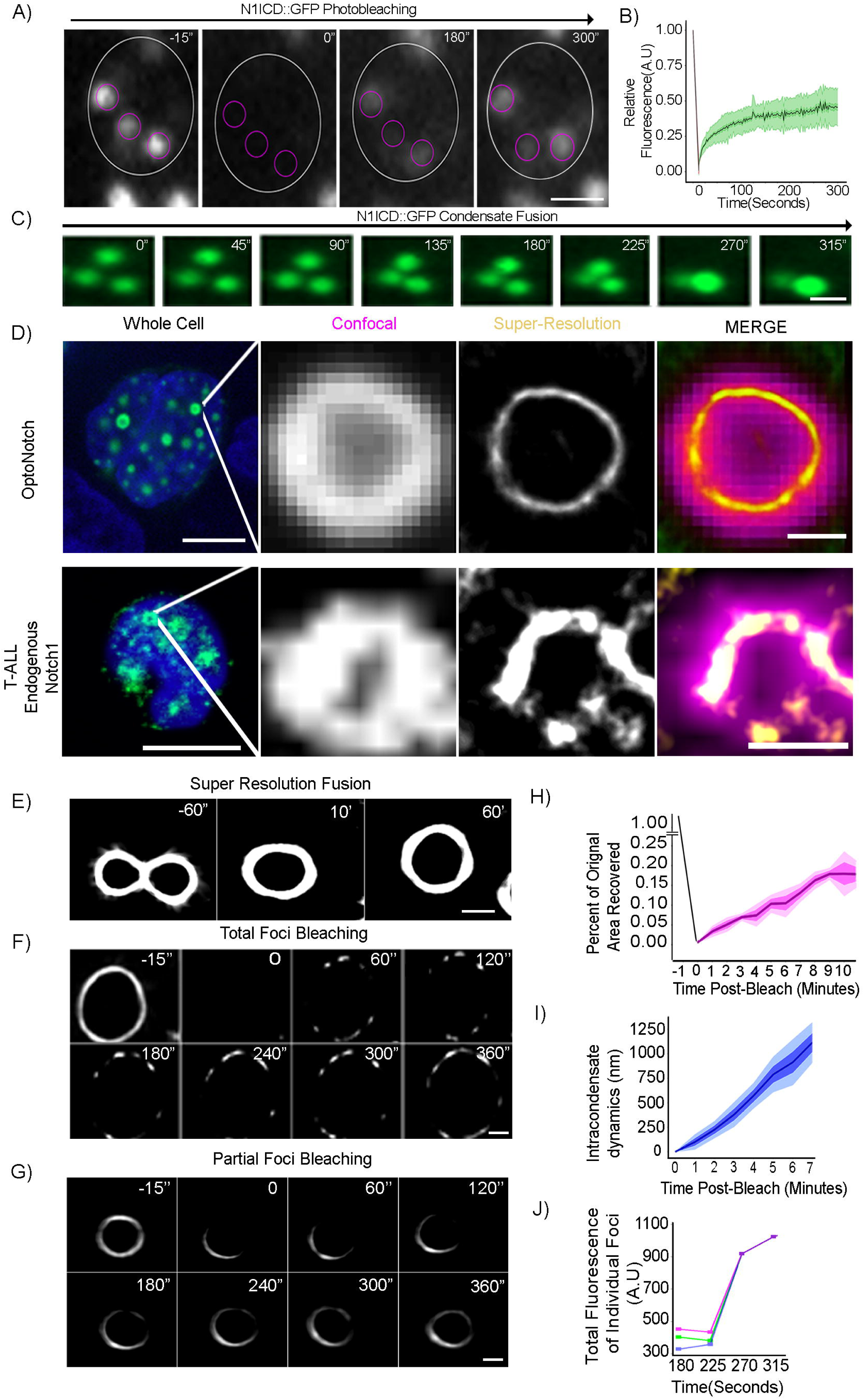
N1ICD Nuclear Foci Exhibit PSCP Properties In Living Cells. A) Photo-bleaching of intranuclear N1ICD foci in OptoNotch-activated HEK293 cells prior to photo-bleaching (left panel; -15”) immediately after photo-bleaching (0”), and at two separate points of recovery (180”, 300”). B) OptoNotch FRAP analysis showing a 46% (+/-14) mobile fraction of the total N1ICD signal with a time-to-half recovery of 24(+/-5) seconds within a single focus over 300 seconds. N = 34 N1ICD photo-bleached foci. C) Example of fusion of N1ICD foci within the nucleus. D) Confocal resolution compared to SRRF imaging resolution of a single OptoNotch N1ICD condensate in a live HEK293 cell (Top) compared to an endogenous N1ICD condensate imaged in a fixed T-ALL cell. Scale bar 10μm (left panel), and 500nm (right panel). E) SRRF imaging of two N1ICD foci undergoing fusion over time. F) Photo-bleaching and subsequent recovery of single N1ICD nuclear foci over time using SRRF imaging. G) SRRF imaging following partial photo-bleaching of a single intranuclear N1ICD focus. H) Percent area recovery following photo-bleaching showing 18+/-7% mobile area. N= 15 N1ICD photo-bleached foci. I) Rate of N1ICD movement within a single focus showing an average rate of movement of 125 nm per minute. N= 11 N1ICD photo-bleached foci. J) Total fluorescence of individual N1ICD foci, (represented by the pink, blue, and green lines) before and after fusion in panel C.

The lack of complete recovery following photo-bleaching suggests that there may exist pools of N1ICD that exhibit differential exchange kinetics. This prompted us to ask whether incomplete recovery is driven by intra-focus heterogeneity, where select sub-domains within individual condensates exhibit differential turnover rates. To address this question, we first performed super-resolution radial fluorescence (SRRF) microscopy to achieve nanometer-scale spatial resolution of either nuclear N1ICD condensates or endogenous Notch1 condensates in T-ALL cells(Figure 3 D, Extended Data Figure 6, Extended Data Figure 7, Movies 4,5). ^61–63^ Our data demonstrate the formation of spherical shell-like intranuclear N1ICD structures, which, when imaged through a single focal plane, present as a ring-like structure. OptoNotch N1ICD and endogenous Notch1 condensates appear to have a non-uniform distribution across the surface of individual condensates, showing potential for entry and exit channels(Figure 3 D, Extended Data Figure 7, Movies 4,5). Using SRRF we were able to visualize condensates undergoing fusion, resulting in an increase in volume with a conservation of Notch signal concentrated in what appeared to be an outer shell surrounding individual condensates, and providing strong evidence of PSCP(Figure 3E).

When subjected to photo-bleaching, N1ICD condensates demonstrate non-uniform recovery across the surface, implying both liquid-like properties and the existence of heterogeneous interactions with unknown factors encapsulated within individual N1ICD condensates(Figure 3F/H, Extended Data Figure 8A). Next, to test the intra-condensate dynamics of individual foci, we photo-bleached only a sub-region of single N1ICD nuclear condensates and observed that a sub-population of molecules within single-condensates exhibit intra-focus movement at an approximate speed of 125 nm/min(Figure 3G/I, Extended Data Figure 8B), demonstrating motility consistent with the formation of intranuclear N1ICD condensates that assemble into spherical, liquid-like shells.

In addition, SRRF imaging revealed clear examples of dynamic changes in the volume (growth and shrinkage) of individual condensates with corresponding fluctuations in N1ICD abundance in the exterior spherical shell(Extended Data Figure 8 C-F, Movie 6). Importantly, we observed the formation and growth of a hollow core in N1CD condensates following initial seeding, demonstrating that condensate shell formation is a function of increases in local N1ICD levels(Extended Data Figure 8G). We next sought to quantify the width of the encapsulating N1ICD shell and compare that to endogenous Notch1 condensates formed in T-ALL cells using SRRF super-resolution imaging and observed no significant difference between endogenous Notch1 and OptoNotch Notch1 condensate shell widths (Extended Data Figure 8H).

Collectively, these results provide strong evidence for the organization of N1ICD into dynamic liquid-like nuclear condensates that possess heterogenous intra- and inter-condensate molecular movement. Importantly, the dynamic growth and shrinkage of intranuclear Notch condensates with interspersed openings suggests exchange of not only N1ICD, but also of transcriptional machineries, nucleotides, and nascent transcripts into-, and out-of the central compartment through anisotropic gaps in the subtending Notch shell.

### N1ICD scaffolds the assembly of functional multiprotein transcriptional condensates

We next sought to determine whether N1ICD foci represent a functional pool of Notch capable of regulating target gene expression.^28–34^ Consistent with functional transcriptional condensates, we observed that N1ICD nuclear condensates consistently colocalize with the canonical Notch protein interactors RBPJ, MAML1, and p300, transcriptional regulators Med1 and BRD4, RNA POLII, as well as nascent mRNA transcripts(Figure 4AB, Extended Data Figure 9 Extended Data Figure 10 A-D). We observed that there is no significant difference in the total percentage of colocalization between N1ICD and any of the co-staining components except for p300, which showed a significantly more variable colocalization coefficient (Figure 4 C). To further investigate the proportional colocalization, we next analyzed the proportion of condensates that were positive for each of the co-staining components as well as the relative proportion of these individual components that are localized within N1ICD condensates. This approach allowed us to quantify the total proportion of Notch1 condensates that contain each of these components, as well as to determine whether Notch1 condensates function to preferentially concentrate any of these factors. We found that there is a significant enrichment in the total amount of MAML1 and RBPJ within N1ICD condensates compared to all other components tested, and that the total amount of Med1 is significantly less than all other components(Figure 4D). We observed, however, that there is a significantly larger population of N1ICD condensates that are positive for MAML1, RBPJ, RNA PolII, and MED1 than those that are positive for BRD4, P300 and BRD-UTP (Figure 4D), which is consistent with the essential roles of RBPJ and MAML1 in N1ICD condensate formation through transactivation complex assembly, and RNA PolII and Med1 in promoting transcriptional activation. Consistent with these results, we also observed that RNA localizes to N1ICD condensates, as demonstrated by co-staining of OptoNotch-activated HEK293 cells with live RNA dye(Extended Data Figure 10 E). In addition, we observed target gene specificity using combination fluorescence protein immunostaining and RNA *in situ hybridization,* which revealed that nascent mRNA transcripts of the Notch target genes Hes1, Hes5, and Hey1 localize precisely to N1ICD condensates(Figure 4E/F/G/H Extended Data Figure 11), whereas sense RNA controls do not exhibit a detectable signal(Figure 4F). Taken together, these results, strongly suggest that N1ICD condensates encapsulate key transcriptional factors, are transcriptionally active, and can facilitate target gene expression over baseline.^3,5,12,17,64^

**Figure 4:**
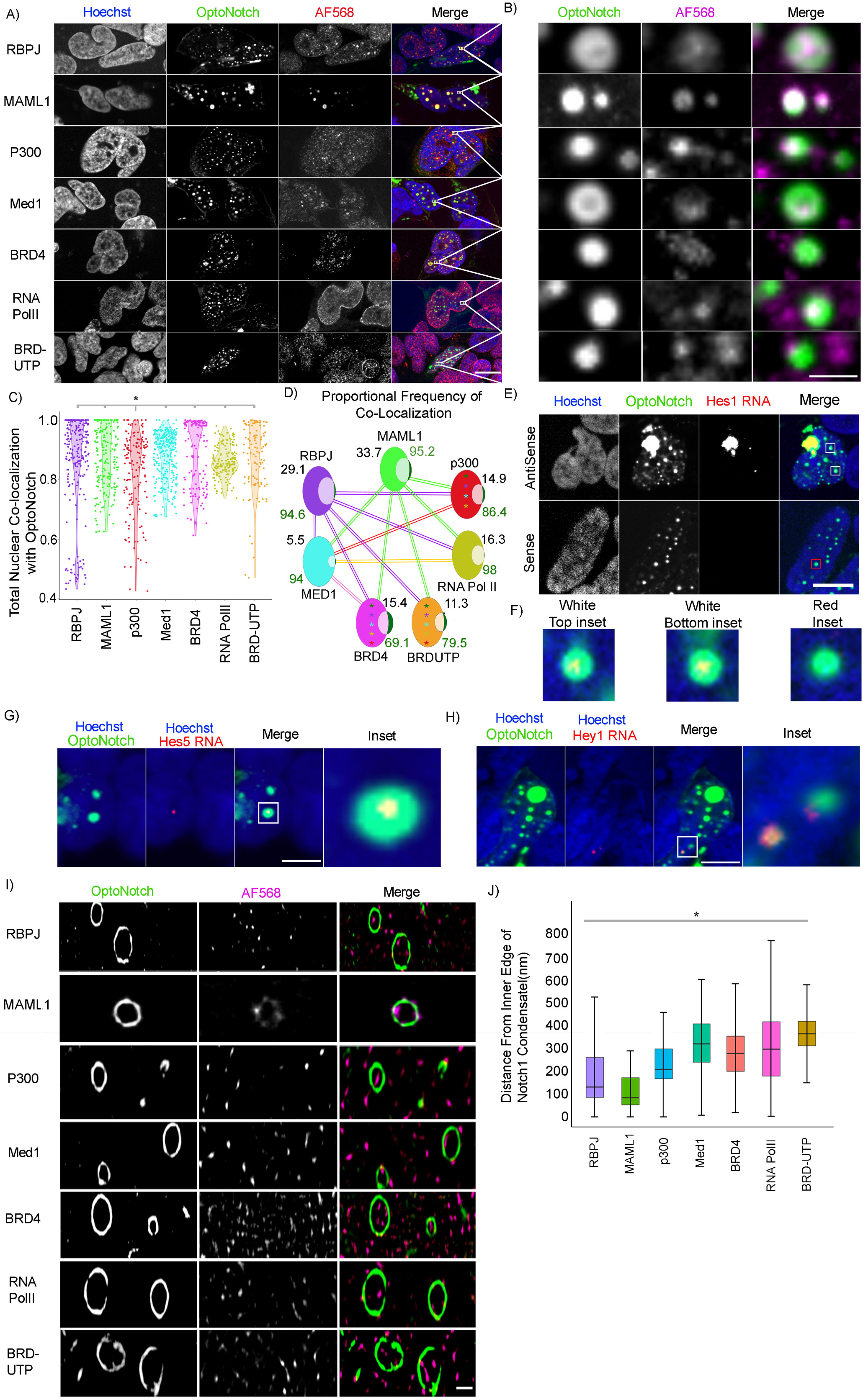
Notch1 Scaffolds The Assembly Of Functional Multiprotein Transcriptional Condensates. A) GSI-treated OptoNotch-activated cells co-stained for Notch1 protein and either RBPJ, MAML1, p300, Med1, BRD4, RNAPOLII, or nascent mRNA transcripts (BRDUTP). B) Inset of individual N1ICD condensates from panel A. C) Total proportion of Nuclear N1ICD that colocalizes with RBPJ, MAML1, p300, Med1, BRD4, RNA PolII, and BRDUTP is 0.862(+/- 0.162), 0.881(+/-0.0938), 0.826(+/-0.132), 0.881(+/-0.0788), 0.885(+/-0.114), 0.862(+/-0.0517), and 0.875(+/-0.112) respectively. N=100 nuclei per condition. *p<0.01 One-way ANOVA+Tukey post-hoc. D) Venn diagrams of the proportional number of nuclear N1ICD condensates that contain RBPJ, MAML1, p300, Med1, BRD4, RNA PolII, and BRDUTP, respectively, is 0.946(+/- 0.0858), 0.957(+/-0.0363), 0.864(+/-0.199), 0.940(+/-0.0634), 0.690(+/-0.273), 0.987(+/-0.0658), and 0.794(+/-0.185) (Green number), and the total amount of RBPJ, MAML1, p300, Med1, BRD4, RNA PolII, or BRDUTP that is contained within N1ICD condensates with respect to the total cellular content is 0.291(+/-0.122), 0.337(+/-131), 0.149(+/-0.093), 0.055(+/-0.015), 0.154(+/-0.062), 0.163(+/-0.121), and 0.113(+/-0.076) (Black number). Lines represent p<0.01 One-way ANOVA+Tukey post-hoc comparisons where the colour denotes which component is significantly greater and the connection shows the comparison (i.e., the green line connecting BRD4 and MAML1 shows that there is a significantly higher proportion of the total amount of MAML1 inside of N1ICD condensates compared to BRD4. * p<0.01 One-way ANOVA+Tukey post-hoc where the colour denotes which component is significantly greater and bubble it is within represents the comparisons (i.e., the green asterisk in BRD4 represents that MAML1 is found in significantly more N1ICD condensates. N= 100 nuclei per condition. E) Hes1 antisense (top) and sense (bottom) RNA *in-situ* in GSI treated, OptoNotch activated, HEK293 cells. F) Insets from Panel E Showing nascent Hes1 expression inside of N1ICD condensates (left and centre) and a Condensate showing no overlap in the sense control(right). G) RNA *in situ* hybridization against Hes5 in HEK293 cells showing complete colocalization of nascent Hes5 transcriptional foci with nuclear N1ICD condensates. H) RNA *in situ* hybridization against Hey1 in HEK293 cells showing complete colocalization of nascent Hey1 transcriptional foci with nuclear N1ICD condensates. I) SRRF images of N1ICD colocalization with RBPJ, MAML1, P300, Med1, BRD4, RNAPolII, or nascent mRNA (BRD-UTP) in cells from panel A. J) Measurements of individual foci from panel I quantifying the distance from the centre of any focus of either RBPJ, MAML1, p300, Med1, BRD4, RNA PolII, or BRD-UTP fluorescence to the nearest inner edge of the N1ICD condensate within which it is contained. Distances equal 187(+/-135.5), 114(+/-76.7), 228(+/-101.7), 317(+/-120.6), 276(+/-117), 319(+/-173), and 362(+/- 97) nm, respectively. *p<0.01 One-way ANOVA+Tukey post-hoc. N= 2000 condensates measured per condition over 50 cells. A/E/G/H Scale bar 10μm. B scale bar 2 μm. I Scale bar 500nm. All data acquired in HEK293. See Source Data Figure 4

Next, we sought to quantify the relative distance between core Notch transcriptional regulators of interest and the edge of the N1ICD condensate shell using SRRF microscopy(Figure 4I/ Extended Data Figure 12). To do so, we measured the total distance from the core of co-staining fluorescence to the inner edge of the N1ICD condensate signal(Figure 4J). We observed that each component is localized to partially overlapping regions within N1ICD condensates(Figure 4J). Specifically, we observed a spatially ordered distribution of protein enrichment where MAML1 showed the closest proximity to the Notch1 shell, followed by RBPJ, p300, Med1, BRD4, and nascently transcribed RNA residing largely in the centre of N1ICD condensates, and thus most distant from the shell, with RNA POLII appearing to have the most diffuse and variable distribution from the edge of the N1ICD condensate shell (Figure 4J). Collectively, these results demonstrate that N1ICD encapsulates and enriches transcriptional components essential for Notch target gene expression inside of transcriptional condensates, thereby facilitating the expression of known Notch1 target genes. Importantly, this is true even under pharmacological inhibition of endogenous Notch signalling, conclusively demonstrating the functionality of OptoNotch.

### Quantification of the transcriptional activity of N1ICD transcriptional condensates

Based on our previous observation that OptoNotch can activate Hes1 expression in a light-dependent manner, we sought to investigate the relationship between OptoNotch transcriptional foci and the frequency, amplitude, and duration of Notch1 target gene activation using a novel Hes1-Live-RNA system that we developed(Figure 5A). ^64,65^ To do so, we first benchmarked our Hes1-Live-RNA reporter by transiently transfecting our MS2/MCP-based system into cells co-expressing OptoNotch, we visualized the spatial distribution of OptoNotch foci with respect to sites of nascent nuclear Hes1-Live-RNA transcription, and compared this distribution to that of a transcriptionally active, Notch-insensitive promoter; EF1α(Figure 5B/C). Consistent with the role of N1ICD in activating Hes1, but not EF1α transcription, we observed a high degree of colocalization between nascent Hes1 RNA reporter transcriptional foci and N1ICD(Figure 5B,5C-Top). We quantified this colocalization by measuring the total distance between any given Hes1-Live RNA or EF1α-Live RNA focus to the nearest N1ICD condensate and observed a significant increase in distance between N1ICD condensates and EF1α-Live RNA foci in comparison to Hes1-Live RNA foci(Figure 5D), implying specificity of N1ICD condensates in activating the Hes1 reporter. In addition to this, we observe that 73.6(+/-15%) of all OptoNotch condensates, in dually transfected cells, show some level of Hes1-live-RNA signal, where 92%(+/-6%) percent of nuclear Hes1-Live-RNA foci co-localize to OptoNotch condensates. In contrast, 0% of OptoNotch condensates are found to be co-localizing with Ef1Alpha-Live-RNA foci. Consistent with a concentration-dependent relationship between N1ICD abundance and Hes1 transcriptional output in Notch1 transcriptional condensates, we observed a direct, linear relationship between the relative fluorescence of N1ICD and Hes1-Live-RNA transcriptional foci(Figure 5E). Similar to our transient transfection data, stable Hes1-Live-RNA cells showed transcriptional activity under endogenous Notch signalling levels, which decreased in response to GSI treatment(Extended Data Figure 13 A/B). Considering that OptoNotch activation is orthogonal to, and independent from endogenous Notch activity, we observed successful rescue of activity following OptoNotch activation under simultaneous blockade of endogenous Notch signalling through GSI inhibition(Extended Data Figure 13). Using stably transfected Hes1-Live-RNA cells, we observed numerous examples of a temporal correlation between the formation of individual nuclear N1ICD condensates, proceeded closely in time by the appearance of Hes1-Live-RNA foci in their centre(Figure 5F). Importantly, we also observed multiple instances of fusion between transcriptionally active condensates, suggesting that individual N1ICD foci encapsulate multiple distinct genomic loci, and can thereby potentially regulate multiple target genes simultaneously (Figure 5G). Moreover, we observed a time-dependent correlation between the intensity of N1ICD transcriptional condensates and Hes1-Live-RNA fluorescence intensity in instances of colocalization between the two signals(Figure 5H/I/J). In addition, we observed nuclear N1ICD transcriptional condensates also exhibited an ability to increase the duration of Hes1-Live-RNA activity at nascent transcriptional foci when Hes1-Live-RNA activity is localized to an N1ICD transcriptional condensate(Figure 5K). Lastly, the Hes1-Live-RNA foci that formed in response to OptoNotch activation were significantly brighter than those formed within control cells lacking OptoNotch, suggesting an increase in transcriptional output (Figure 5L). Taken together, these data further suggest that N1ICD transcriptional foci are functional in driving target gene expression and that Notch1 target gene expression is directly proportional to the total abundance of N1ICD within a transcriptional condensate, demonstrating a direct relationship between Notch1 abundance and transcriptional output. Collectively, these data strongly suggest that increases in intranuclear N1ICD levels increases the size of individual N1ICD transcriptional condensates, which, in turn, increase both the duration and intensity of target gene transcription, providing further evidence for a functional role for Notch1 transcriptional condensates.

**Figure 5:**
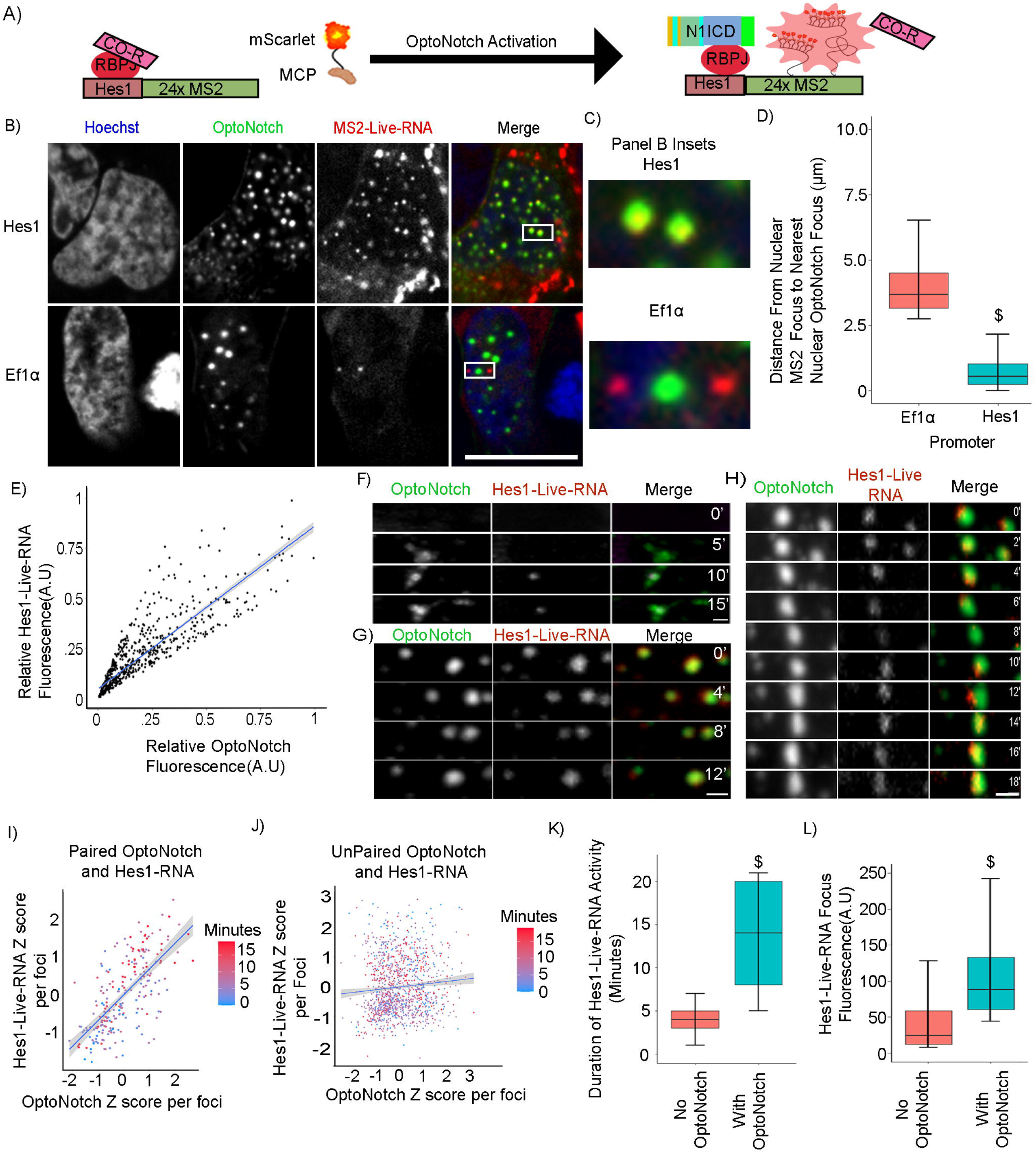
Analysis of N1ICD condensate transcriptional bursting dynamics. A) Schematic representation of the Hes1-Live-RNA reporter construct architecture and function. B) Hes1-Live-RNA reporter(top) or Ef1α-Live-RNA control(bottom) expression in OptoNotch-activated HEK293 cells. Scale bar 10μm. C) Insets from panel B from either Hes1-Live-RNA (top) or Ef1α-Live-RNA (bottom). D) Average distance between the centre of a Live-RNA nascent transcriptional focus and the nearest N1ICD transcriptional condensate seeing an average distance of 0.693μm (+/-0.525) with the Hes1 promoter, or 4.11μm (+/-1.39) for Ef1α promoter. N=7000 foci per condition. E) Pearson correlation of Hes1-Live-RNA nascent transcriptional foci and N1ICD fluorescence in OptoNotch-activated HEK293 cells showing colocalization with an R^2 of 0.877. N=800 condensates measured. F) OptoNotch activation facilitating expression of Hes1-Live-RNA inside of an individual Notch1 condensate. G) Hes1-Live RNA containing intranuclear N1ICD condensates exhibit fusion in HEK293 cells without loss of Hes1-Live-RNA. H) Colocalization between N1ICD and Hes1-Live-RNA fluorescence reveals fluctuations in Hes1-Live-RNA abundance over time following OptoNotch activation in HEK293 cells. I) Correlation Z-scores of the relative intensity of each channel in relation to time post OptoNotch activation in foci containing both N1ICD and Hes1-Live-RNA. Blue line represents Pearson correlation with an R^2 of 0.684. N = 18 foci. J) Correlation Z-scores of the relative intensity of each channel in relation to time post OptoNotch activation in foci that contain only N1ICD or Hes1-Live-RNA. Blue line represents a Pearson correlation with an R^2 of 0.09. N=45 foci. K) Quantification of the duration of Hes1-Live-RNA transcriptional bursting shows Hes1-Live-RNA transcriptional bursts last for an average of 3.9(+/-0.7) minutes in wild-type cells, whereas Hes1-Live-RNA transcriptional foci that colocalize with intranuclear N1ICD transcriptional condensates exhibit an increased bursting time with an average of 18.9(+/-4.2) minutes. N=1200 foci per condition. L) Comparison of the fluorescence intensity of Hes1-Live-RNA fluorescence either not associated with a N1ICD condensate with an average intensity of 44.8(+/-48.5) or when associated with a N1ICD condensate having an average intensity of 104(+/-52.5). N=1200 foci per condition $ p<0.01 on student T-test. *p<0.01 one-way ANOVA+Tukey post-hoc. F/G/H Scale bar 2 μm. All data acquired in HEK293. See Source Data Figure 5

### Investigating the role of Notch transcriptional activation complex assembly on N1ICD condensate formation

To further identify the factors responsible for Notch transcriptional condensate formation, we investigated the role of Notch1 transcriptional activation complex assembly in Notch1 PSCP either *via* knockout of RBPJ or pharmacological disruption of the N1ICD-RBPJ complex. In contrast to wild type and DMSO-treated cells, where OptoNotch activation resulted in the formation of prominent and abundant intranuclear foci, we observed a significant reduction in the number of N1ICD transcriptional condensates in RBPJ knockout cells, and when Notch transcription activation complex assembly was disrupted using CB-103; a potent and specific disruptor of Notch transcriptional complex assembly(Figure 6A/B).^10,16^ Specifically, we observed that RBPJ KO cells showed a significant reduction in nuclear OptoNotch N1ICD levels that was not observed in CB-103 treated cells(Figure 6C/D). These results suggest that transcriptional condensate formation is stabilized and promoted through RBPJ-mediated N1ICD anchoring, however, the ankyrin repeat domain-mediated anchoring is not required for Notch condensate formation as we observed N1ICD condensates in the absence of RBPJ (Figure 6C/D).

**Figure 6:**
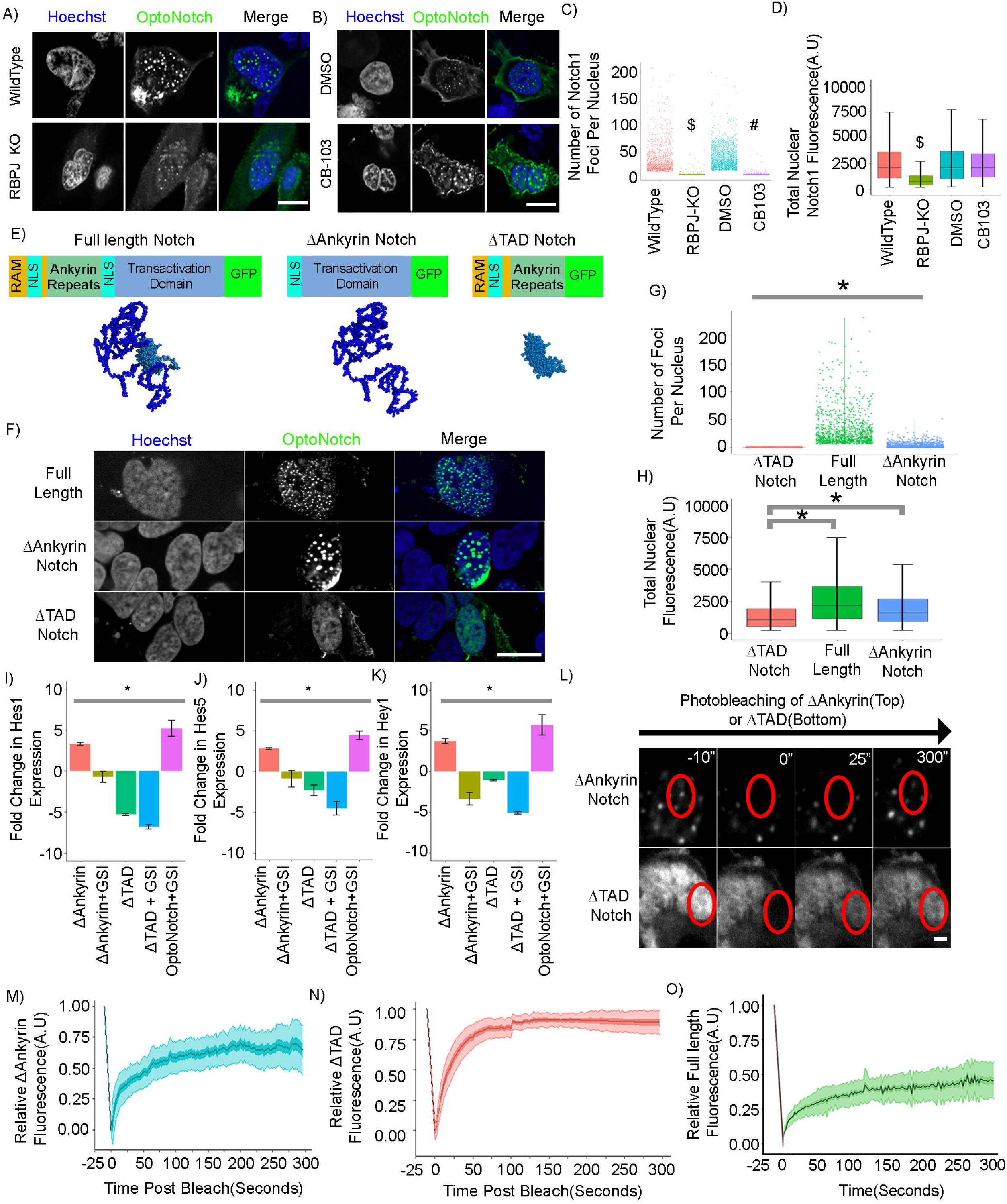
Structure/Function Analysis Of The N1ICD With Respect To N1ICD Nuclear Transcriptional Condensate Formation. A) Activated OptoNotch in wildtype and RBPJ knockout HeLa cells. B) Activated OptoNotch in DMSO-treated, and CB-103-treated wild-type HEK293 cells. C) Quantification of the number of N1ICD condensates per nucleus in A and B, showing that Wildtype, RBPJ KO, DMSO-treated wildtype, and CB-103-treated wildtype cells contain an average of 30.53 (+/-30.14), 0.9(+/-4.26), 32.016(+/-21.99), and 0.54(+/-0.44) N1ICD condensates per nucleus, respectively. N= 700 nuclei per condition. D) Total nuclear fluorescence from panel A and B showing that Wildtype, RBPJ KO, DMSO-treated wildtype, and CB-103-treated wildtype cells contain an average fluorescence intensity of 2581(+/-2020), 1376(+/-2283), 2692(+/-2514), and 2849(+/-2515) AU of N1ICD. N= 700 nuclei per condition. E) Structural schematic of the full-length, ΔAnkyrin, and ΔTAD OptoNotch constructs (top) with corresponding AlphaFold structure predictions(bottom). Full length, ΔAnkyrin, and ΔTAD OptoNotch after light activation in HEK293 cells. G) Quantification of the number of foci per nucleus in F. ΔTAD Notch-expressing cells contain zero foci per nucleus, in comparison to full length Notch- and ΔAnkyrin Notch-expressing cells, which contain 30.8(+/-31.3), and 2.5(+/-4.09) foci per nucleus, respectively. N=700 nuclei per condition. H) Quantification of the total nuclear fluorescence intensity from F. ΔTAD, full length, and ΔAnkyrin Notch-expressing cells shows 1631(+/-2561), 2692(+/-2927), and 2670(+/-3859) AU of GFP signal, respectively. N= 700 cells measured per condition. I) Quantification of Hes1 expression by qPCR in activated OptoNotch-expressing cells with either ΔAnkyrin, ΔAnkyrin+GSI, ΔTAD, ΔTAD+GSI, or full-length + GSI show a fold change in Hes1 expression of 3.307(+/-0.157), -0.740(+/-0.670), -5.316(+/-0.122), -6.850(+/-0.278), and 5.200(+/-1.001), respectively. J) Quantification of Hes5 expression by qPCR in activated OptoNotch-expressing cells with either ΔAnkyrin, ΔAnkyrin+GSI, ΔTAD, ΔTAD+GSI, or full-length + GSI show a fold change in Hes5 expression of 2.848(+/-0.0956), -0.898(+/-1.003), -2.303(+/-0.643), -4.527(+/- 0.853), and 4.446(+/-0.504), respectively. K) Quantification of Hey1 expression by qPCR in activated OptoNotch-expressing cells with either ΔAnkyrin, ΔAnkyrin+GSI, ΔTAD, ΔTAD+GSI, or full-length + GSI show a fold change in Hey1 expression of 3.706(+/-0.300), -3.370(+/-0.769), -1.069(+/-0.096), -5.110(+/-0.149), and 5.697(+/-1.246) respectively. L) Fluorescence photo-bleaching recovery experiments of nuclear ΔAnkyrin and ΔTAD OptoNotch. Photo-bleached regions are shown as red circles in each experiment. Scale bar 2um. M) Fluorescence photo-bleaching recovery of nuclear ΔAnkyrin showing a mobile fraction of 72 (+/-17) % with a half-recovery time of 22(+/-5) seconds. N=40 photo-bleaching experiments. Central line -mean, dark green-SEM, light green-SD. N) Fluorescence photo-bleaching recovery of nuclear ΔTAD showing a mobile fraction of 95% (+/-4) with 14(+/-2) seconds to half recovery. N=40 photo-bleaching experiments. Central line -mean, dark red-SEM, light red-SD. O) Replicate of data from figure 3B to allow for visual comparison of bleach kinetics of all three variants of OptoNotch. $p<0.01 student t-test comparing RBPJ KO to Wildtype. #p<0.01 student t-test comparing CB-103 to Vehicle. *p<0.01 One-way ANOVA+Tukey post-hoc. A/B/F scale bar = 10um. See Source Data Figure 6

Next, we assessed the necessity of the N1ICD:RBPJ interaction for transcriptional condensate formation by generating and studying two additional GFP-tagged OptoNotch alleles; Δ-Ankyrin N1ICD and Δ-TAD N1ICD(Figure 6E). Consistent with our transcriptional activation complex disruption experiments, cells expressing Δ-TAD OptoNotch failed to form intranuclear transcriptional condensates, with cells exhibiting only a diffuse nuclear localization of N1ICDΔTAD(Figure 6F/G/H). In contrast, and consistent with our results both in RBPJ KO cells and with pharmacological disruption of the Notch transcriptional activation complex, ΔAnkyrin OptoNotch-activated cells exhibited a significant decrease in the number of Notch condensates compared to full-length OptoNotch, with only a small, persistent population of condensates (Figure 6G/H). Specifically, we observed that the total nuclear abundance of ΔTAD N1ICD is significantly lower than that of either Δ-Ankyrin N1ICD, or full-length N1ICD, with no observable difference in the total abundance of Δ-Ankyrin N1ICD compared to full-length N1ICD, implying that condensate formation may stabilize the N1ICD against clearance and turnover (Figure 6H). Additionally, ΔAnkyrin-OptoNotch is able to form hollow core condensates alluding to that these condensates are also multi-protein condensates(Extended Data Figure 14). We next explored whether Δ-Ankyrin N1ICD condensates localize to endogenous Notch1 target loci, where we observed that Δ-Ankyrin N1ICD colocalizes with RBPJ(Extended Data Figure 15).

Colocalization between Δ-Ankyrin N1ICD and RBPJ could be mediated by endogenous N1ICD, which contains intact Ankyrin repeats, indirectly recruiting Δ-Ankyrin N1ICD to Notch transcriptional activation complexes through TAD based interactions with MAML1, or a through recruitment by some unknown factor. Based on our Δ-Ankyrin N1ICD colocalization experiments, which showed a greater degree of colocalization with MAML1 compared to RBPJ, we propose that the formation of Δ-Ankyrin N1ICD condensates may arise through interactions between the TAD and MAML1 (Extended Data Figure 15). To test whether the formation of condensates at endogenous Notch target genes can facilitate target gene expression we performed qPCR on cells activated with either Δ-Ankyrin N1ICD, Δ-TAD N1ICD, or full-length N1ICD. We observed that in the presence of endogenous Notch1, Δ-Ankyrin N1ICD can significantly increase Notch target gene expression(Figure 6I-K), and that this effect is completely abolished upon inhibition of endogenous Notch activation(Figure 6I-K). Conversely, when cells are activated with Δ-TAD N1ICD, we observed inhibition of Notch target gene expression (Figure 6I-K). Taken together, our results demonstrate that the N1ICD TAD domain assembles into condensates and is sufficient to facilitate target gene expression when recruited by endogenous full-length N1ICD to active transcriptional condensates. In addition, these data provide evidence that the interaction between the N1ICD and RBPJ is not capable of driving high levels of target gene expression in the absence of the N1ICD TAD domain, and that facilitating the formation of N1ICD condensates increases Notch target gene expression in a concentration-dependent manner.

Considering both the inability of ΔAnkyrin N1ICD to bind to RBPJ and the lack of condensate formation with ΔTAD N1ICD, we next sought to determine whether these two truncated alleles exhibit differential intranuclear mobilities compared to full-length N1ICD. FRAP experiments with ΔTAD N1ICD showed a significantly larger mobile fraction, and a significantly shorter half-recovery time compared to full-length N1ICD(Figure 6L/M/N/O, Movie 7). In addition, we observed a significantly lower mobile fraction of ΔAnkyrin N1ICD, compared to ΔTAD N1ICD, but a significantly higher mobile fraction compared to full-length N1ICD (Figure 6L/M/N/O, Movie 8). In addition, ΔAnkyrin N1ICD exhibited a significantly slower recovery time than ΔTAD N1ICD but showed no significant difference compared to full-length N1ICD(Figure 6L/M/N/O). Taken together, these data demonstrate that Δ-Ankyrin N1ICD exhibits increased intra-condensate mobility in comparison to full-length N1ICD and that the exchange of N1ICD protein in both conditions is regulated by similar interactions, likely mediated by the N1ICD IDR, as we observed no difference in the half-recovery times. Overall, these data provide evidence that anchoring through the RBPJ/ankyrin domain interaction decreases intra-condensate motility, and that mobile fractions of N1ICD condensates are not regulated by canonical RBPJ/N1ICD interactions, but rather through interactions driven by the C-terminal TAD domain.

### N1ICD Transcriptional Condensates Promote Super-Enhancer Looping at the Human *MYC* Locus

To further investigate the role of Notch1 transcriptional condensate assembly in facilitating target gene expression, we employed an established model of Notch-dependent *MYC* proto-oncogene super-enhancer looping.^7–9,51,52,66,67^ It is still unclear, however, how Notch functions to reorganize these distal regions, allowing for a direct effect on MYC expression following Notch binding >1.7Mb away, a distance that is sufficient for measuring promoter/super-enhancer contacts using confocal/super-resolution microscopy (Figure 7A/B).^39^

**Figure 7:**
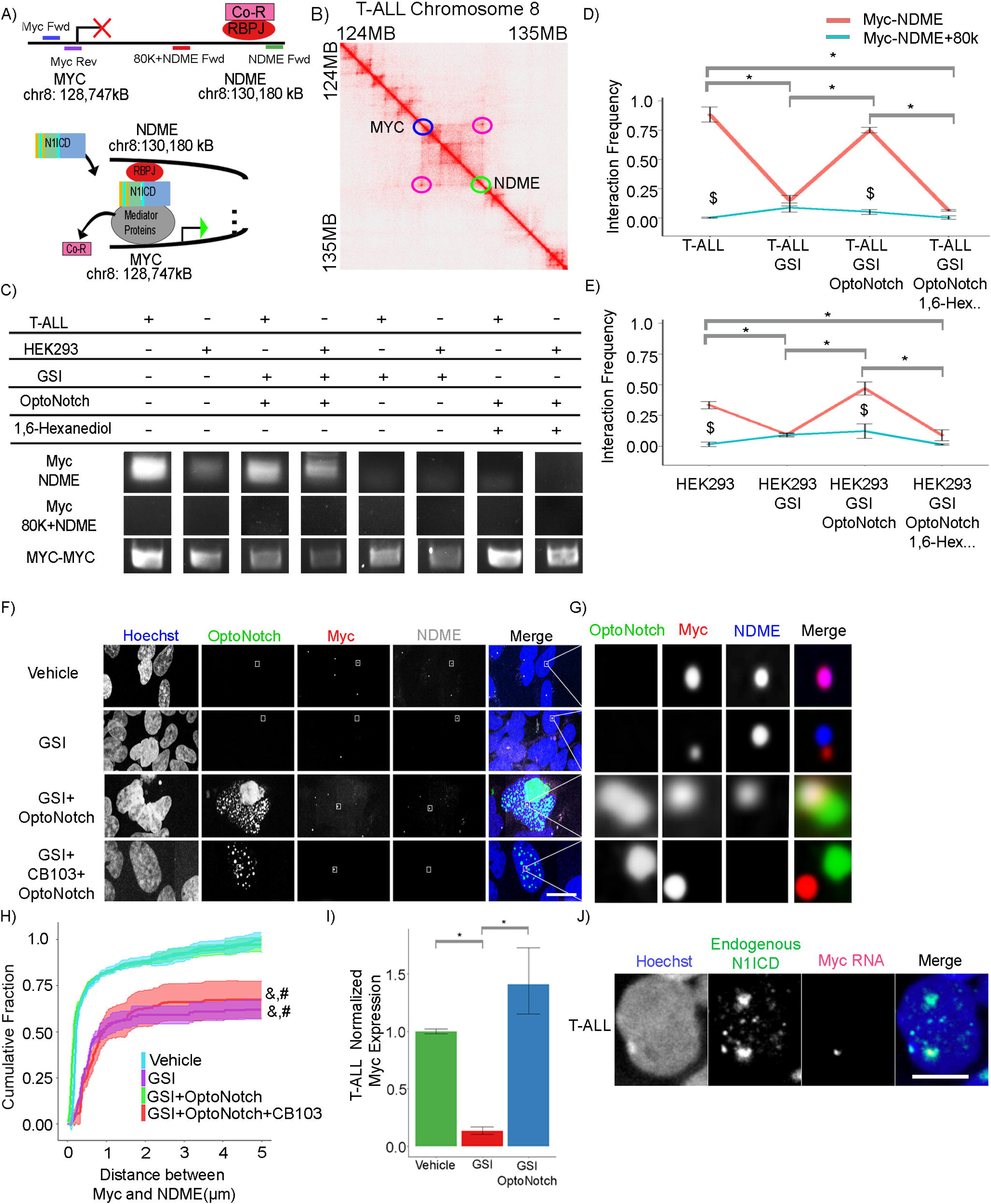
Notch1 Transcriptional Condensates Facilitates Super-Enhancer Looping At The Human MYC Locus. A) Schematic of the Human Myc genomic locus on chromosome 8, containing the MYC and NDME loci, and illustrating a model of Notch-dependent super-enhancer looping. MYC-MYC primers are shown in blue and purple, MYC–NDME primers are shown in blue and green, and the MYC-MYC -NDME+80K are shown in blue and red. Genomic positions are based on GRCh38. B) Hi-C heat map of the Myc locus on chromosome 8 in T-ALL cells highlighting the MYC promoter(Blue circle) and the NDME (Green circle) and the loop that they form in T-ALL cells ( Purple circle). C) Gel electrophoresis of the 3CPCR MYC-NDME, MYC-NDME+80K, and MYC-MYC products from cells treated with vehicle control, GSI, GSI+ N1ICD, or GSI+N1ICD+1,6-hexanediol in either T-ALL or HEK293 cells. D) 3CqPCR Interaction frequency of Myc-NDME(Red) and Myc-NDME+80K(blue) in T-ALL cells from vehicle (0.8833+/- 0.0645[red], 0.0018+/-0.0049[blue]), GSI (0.1476+/-0.0444 [red], 0.0881+/-0.0384[blue]), GSI+OptoNotch(0.7491+/-0.0234[red], 0.0508+/-0.0213[blue]), or GSI+OptoNotch+1,6- Hexanediol(0.0643+/-0.0064[red], 0.024+/-0.0154[blue]) treatments. D) 3CqPCR Interaction frequency of Myc-NDME(Red) and Myc-NDME+80K(blue) in HEK293 cells from either vehicle(0.3336+/-0.0288[red], 0.0149+/-0.0185[blue]), GSI(0.0928+/-0.0092[red], 0.0906+/- 0.0167[blue]), GSI+OptoNotch(0.4683+/-0.0539[red], 0.1212+/-0.0585[blue]), or GSI+OptoNotch+1,6-Hexanediol(0.0872+/-0.0449[red], 0.0118+/-0.0055[blue]) treatments. D/E $p<0.01 student t-test between the MYC-NDME and MYC-NDME+80k within a condition. F) Fluorescence images of DNA-Paint-stained samples of OptoNotch, NDME and the MYC locus in HEK293 cells treated with vehicle, GSI, GSI+N1ICD, and GSI+NICD+CB-103. Scale Bar 10um. G) Insets from corresponding white boxes in panel F. H) Cumulative Fraction of the Distance between the MYC and the NDME loci. N=350 MYC loci per condition. &, # p<0.01 One-way ANOVA+Tukey post-hoc between vehicle or GSI+OptoNotch. I) Quantification of MYC expression by qPCR in T-ALL cells treated with either vehicle, GSI or GSI+ OptoNotch, showing a relative expression of MYC of 1(+/-0.05), 0.133(+/-0.618) and 1.410(+/-0.508), respectively for each condition. J) Myc RNA *in-situ* co-stained for endogenous N1ICD in T-ALL cells. Scale bar 5um. *p<0.01 One-way ANOVA+Tukey post-hoc D/E/I. See Source Data Figure 7

To address this question, we next sought to determine whether N1ICD transcriptional condensates are functional in promoting the assembly of the Notch-dependent *MYC* super-enhancer and concomitant activation of *MYC* gene expression. To demonstrate a dependency upon Notch1 for enhancer looping and the direct interaction between the *MYC* and NDME loci, we first performed 3C-PCR to determine whether the formation of N1ICD transcriptional condensates bring these two distal genomic loci into close physical proximity with one another in both T-ALL cells, in which this interaction has been previously described, and in HEK293 cells, to determine whether MYC super-enhancer looping occurs in this cell type as well(Figure 7C).^68–70^ Following inhibition of Notch signalling, we observed a decrease in the interaction frequency between the *MYC* and NDME loci in comparison to vehicle-treated cells(Figure 7D/E). We subsequently tested whether this effect is rescued by OptoNotch activation under GSI inhibition, and observed a significant increase in the interaction frequency between the *MYC* and NDME loci in comparison to GSI-treated cells alone, with no significant difference in interaction frequency compared to untreated cells(Figure 7D/E). Importantly, we also observed that transcriptional condensate formation is required to recruit NDME to the *MYC* locus, as 1,6-hexanediol treatment resulted in a significant reduction in the interaction frequency between the *MYC* and NDME loci(Figure 7D/E). These effects were seen in both T-ALL and HEK293 cells under each of the various treatment conditions(Figure 7 C-E).

To further confirm this result, we next employed DNA-PAINT to visualize the spatial colocalization of the MYC and NDME loci under different conditions to test the role of Notch condensates in MYC super-enhancer looping(Figure 7F/G). At endogenous levels of Notch activity, we observed that the *MYC* and NDME loci colocalize in prominent intranuclear foci in HEK293 cells. In contrast, following Notch1 inhibition, the *MYC* and NDME loci become spatially distinct, indicating spatial separation of the MYC promoter and NDME locus(Figure 7F/G).^70^ Importantly, we observed that OptoNotch activation in the presence of endogenous Notch1 inhibition results in a rescue of the colocalization between the *MYC* and NDME loci, and that intranuclear *MYC*-NDME loci are localized within intranuclear N1ICD transcriptional condensates (Figure 7F/G). Considering that these data are consistent with a role for the Notch transcriptional activation complex in *MYC*-NDME enhancer looping, we next sought to determine whether specifically disrupting the Notch transcriptional activation complex inhibits MYC super-enhancer looping. We found that CB103 treatment, which blocks Notch transcriptional activation complex assembly, results in a significant increase in the distance between the *MYC* and NDME loci, implying blockade of MYC super-enhancer looping, despite the presence of nuclear N1ICD condensates after OptoNotch activation(Figure 7F/G/H). These results demonstrate that MYC super-enhancer looping is dependent upon assembly of an intact Notch transcriptional activation complex and that N1ICD condensates promote super-enhancer looping in the absence of endogenous Notch signalling as demonstrated by our observation that OptoNotch activation in GSI-treated cells rescues physical association between *MYC* and NDME genomic loci through their recruitment to N1ICD transcriptional condensates (Figure 7H).

Lastly, to demonstrate the ability of OptoNotch to rescue *MYC* expression, and thus provide supporting evidence of MYC super-enhancer looping through functional association between the *MYC* and NDME loci, we next performed qPCR on *MYC* expression in both T-ALL cells, which exhibit a Notch dependency for *MYC* expression, and in HEK293 cells. ^7–9,51,52,66^ Quantitative analysis of MYC expression demonstrated that Notch inhibition results in a concomitant decrease in *MYC* expression in both cell lines, and that OptoNotch can rescue *MYC* expression in both cell lines despite inhibition of endogenous Notch signalling(Figure 7I,Extended Data Figure 16). To demonstrate that endogenous N1ICD condensates are sites of active Myc transcription in T-ALL cells we performed RNA *in situ* hybridization against MYC RNA in untreated T-ALL cells that were co-stained for N1ICD(Figure 7J). We observed nascent MYC transcriptional foci that clearly colocalize with an endogenous N1ICD condensate. Lastly, we quantified MYC expression in HEK293 cells using RNA *in situ* hybridization, where we observed that Notch inhibition through GSI treatment significantly decreased MYC transcript levels, whereas OptoNotch activation significantly increased MYC transcript levels, demonstrating both the necessity and sufficiency of N1ICD in regulating MYC expression in these cells (Extended Data Figure 16).

## Discussion

Here, we provide direct evidence that the intrinsically disordered tail of the human N1ICD facilitates transcriptional activation in a Notch-activity-dependent manner through the assembly of functional transcriptional condensates. Our initial *in vitro* experiments provide evidence that purified N1ICD is capable of undergoing PSCP in a salt- and concentration-dependent manner. By employing GSI’s to block Notch signalling in constitutively active T-ALL cells, and surface-immobilized DeltaMAX to activate endogenous Notch signalling in HEK293 cells we show that endogenous N1ICD undergoes PSCP and that both the abundance and size of intranuclear N1ICD condensates are regulated by the level of Notch activity. To investigate this relationship, we developed a novel Optogenetic tool, OptoNotch, that possesses light-dependent, gamma secretase cleavage-independent activity, and which affords precise spatial and temporal control over intranuclear levels of GFP-tagged N1ICD variants, and concomitant transcriptional activation of Notch1 target genes.^53,57^ The results of our titration experiments using OptoNotch demonstrate a direct relationship between nuclear N1ICD abundance, intranuclear condensate formation, and target gene expression. We also show that OptoNotch-activated full-length N1ICD spontaneously self-assembles into highly dynamic intranuclear spherical condensates that exhibit seeding, growth, and shrinkage over time. Importantly, we demonstrated that the OptoNotch system functions within the dynamic range of endogenous Notch1 signalling capacity through direct comparison with endogenous intranuclear N1ICD condensates formed in response to DeltaMAX-mediated activation and in a relevant T-ALL model cell line.

To further investigate condensate dynamics and morphology, we subsequently employed SRRF microscopy, and show that N1ICD spontaneously self-assembles into highly dynamic intranuclear spherical shells, similar to what is seen with endogenous Notch1 in T-ALL cells. Notch1 condensates exhibit exclusion of N1ICD from a central core, and, when subjected to photo-bleaching, demonstrate non-uniform recovery, implying both liquid-like properties and the existence of heterogeneous interactions with unknown factors encapsulated within individual N1ICD condensates. Our observation that N1ICD condensates exhibit dynamic motility and anisotropic recovery provides evidence for liquid-like properties of the N1ICD condensate shell, consistent with PSCP.

Importantly, we provide examples in which individual N1ICD condensates exhibit dynamic fluctuations in N1ICD abundance and in condensate volume, implying that these structures are indeed highly dynamic and have liquid-like properties. Importantly, our observation of the formation and growth of a hollow core in N1ICD condensates following initial seeding demonstrates that condensate shell formation is coupled to condensate growth through a process consistent with Ostwald ripening. Importantly, our observation of N1ICD condensate fusion, during which both N1ICD levels and total condensate volume are conserved, provides further demonstration of liquid-like properties of the shell, and strong evidence for PSCP.

Using multicolour fluorescence immunohistochemistry, we show that N1ICD transcriptional condensates increase target gene expression by encapsulating, and thereby enriching the core transcriptional activation complex interactors RBPJ and MAML1, and to a lesser extent, key transcriptional regulators and machineries, including Med1, P300, BRD4, and RNA Polymerase 2, which we precisely mapped to distinct regions within N1ICD condensates. ^62,63^ In accordance with this finding, we demonstrate that N1ICD condensates are transcriptionally active and that canonical Notch1 target genes Hes1, Hes5, and Hey1, are spatially localized to the centre of N1ICD condensates, suggesting that N1ICD assembles into active transcriptional condensates. To further test the input-output relationship between Notch1 condensate formation and transcriptional activity, we developed and employed a live Hes1 transcriptional reporter, to visualize and quantify Hes1 transcriptional activity in living cells and show that Hes1 transcription is directly proportional to the level of N1ICD in individual condensates. In addition, we show that transcriptional burst duration and intensity, rather than burst frequency, are extended and increased in proportion to the intranuclear abundance of N1ICD and its assembly into transcriptional condensates.

We then determined that the TAD domain alone is sufficient for intranuclear condensate assembly and that it is capable of facilitating Notch1 target gene expression when a ‘seed’ of endogenous N1ICD is present, but that it is incapable of activating target gene expression on its own in the absence of the RAM domain and ankyrin repeats, which are responsible for its physical interaction with RBPJ^5,16–18^. Conversely, we show that the N1ICD ankyrin repeats alone are incapable of forming intranuclear Notch1 condensates, or increasing target gene expression. Taken together these data demonstrate the necessity of the TAD domain in condensate assembly and highlight its importance in facilitating target gene expression.

Lastly, we employed an established model of Notch-dependent *MYC* proto-oncogene super-enhancer looping to investigate the role of N1ICD transcriptional condensates in super-enhancer assembly at the Human MYC locus. Using a combination of 3C-qPCR and DNA-Paint we show a dependency upon endogenous Notch activity, and a sufficiency of ectopic N1ICD, in promoting NDME-dependent MYC super-enhancer looping, whereby 1,6-hexanediol-soluble N1ICD transcriptional condensates increase the interaction frequency between these two distal genomic loci. These results demonstrate both the necessity and sufficiency of N1ICD for NDME-dependent MYC super-enhancer looping. We further show that dissolution of the Notch transcriptional activation complex through CB-103 treatment result in the separation of the MYC and NDME loci without dissolution of N1ICD condensates, demonstrating that anchoring of the N1ICD by the transcriptional activation complex is essential for MYC superenhancer looping. These results provide direct evidence that *MYC* and NDME genomic loci colocalize in intranuclear N1ICD transcriptional condensates, and that their colocalization depends upon formation of an intact Notch1 transcriptional activation complex, suggesting a role for N1ICD condensates in mediating MYC enhancer/promoter interactions. This hypothesis is further supported by our observations that endogenous Notch activity is necessary for MYC expression and that OptoNotch activation increases MYC expression as quantified by qPCR and RNA *in situ* hybridization in both T-ALL and HEK293 cells. Collectively, our results provide evidence that PSCP-driven Notch1 transcriptional condensate formation represents a novel mechanism through which Notch signalling facilitates the assembly and activation of the *MYC* super-enhancer.

Previous research has demonstrated that transcriptional condensates regulate gene expression through a non-equilibrium process that provides dynamic feedback through its RNA product, supporting a model where RNA abundance provides positive and negative feedback on transcription via regulation of electrostatic interactions.^71,72^ Our observation that N1ICD spontaneously self-organizes into heterogeneous spherical shells with interspersed Notch-free regions, suggests the presence of entry/exit channels for transcriptional components.

Considering that Notch1 transcriptional condensates exhibit dynamic growth and reduction phases in terms of both volume and N1ICD abundance, that transcriptionally active condensates are capable of homotypic fusion while retaining both N1CD and target gene transcripts, and that N1ICD condensates do not dissolve in response to transcriptional bursting, we propose a model in which N1ICD condensates allow for transit of transcriptional regulatory machinery, nucleotide substrates, and transgenic proteins into-, and nascent RNA transcripts out- of, Notch transcriptional condensates. These NICD-free channels may allow for the alleviation of electrostatic repulsion driven by RNA transcript accumulation inside individual N1ICD condensates, thereby increasing transcriptional burst duration by reducing the frequency of condensate dissolution.

Recent work has demonstrated that genome topology is a critical feature of gene control and that transcriptional condensates provide an important regulatory layer to the three-dimensional organization of the genome.^73,74^ Our observation of fusion between Notch transcriptional condensates with retention of nascent Hes1 transcripts implies the coalescence of multiple distinct genomic loci into single transcriptional condensates. Future studies that integrate the methods we have employed in combination with strategies capable of providing information about dynamic genomic landscapes, on a single cell level, will allow for a deeper understanding of the mechanisms that drive Notch-mediated transcriptional regulation. In addition, further investigation using super-resolution microscopy and orthogonal fluorescent labelling of relevant proteins should be aimed at addressing the dynamic flux of transcriptional machineries (i.e. Med1, RNA Polymerase, etc.) and nascent transcripts into and out of Notch transcriptional condensates. This approach would allow for further characterization of the mechanism(s) through which Notch increases transcriptional burst duration by physically stabilizing the transcriptional machinery, while simultaneously alleviating the destabilizing effects of nascent transcripts by allowing for their unimpeded efflux.

As an extension of this work and considering the large number of computationally predicted post-translational modification sites identified in the N1ICD (https://elm.eu.org), we anticipate that there exists a vast unexplored landscape of Notch proteoforms that modulate transcriptional condensate dynamics and function. Each proteoform may represent a variant that has integrated multiple layers of cellular signalling inputs in distinct ways and may thus be capable of uniquely modulating the transcriptional output of discrete target genes in distinct ways. In future, further exploration of the dynamics and function of Notch IDR’s involving human and non-human NICDs, many of which differ in the presence and/or length of IDRs, will help to shed light on the molecular ‘grammar’ of IDR function in Notch signalling by characterizing how discrete changes in IDR’s influence their ability to undergo PSCP and to spontaneously self-assemble into active transcriptional condensates.^1,11^

## METHODS

### Molecular Cloning

OptoNotch constructs (OptoNotch (Full-length N1ICD [aa1779-2555]), OptoNotch^mut^ (mutant TEV cleavage sequence), OptoNotchΔTAD [aa1779-2169], OptoNotchΔAnkyrin [aa2170-2555], Cry2-cTEV) were generated by PCR and Gibson assembly to be subsequently sub-cloned into a modified MXS chaining vector containing a CMV promoter and BGHpa Tail. ^75^

OptoNotch comprises two separate proteins: one containing Cry2PHR, a protein that, when illuminated with blue light, will interact with its binding partner CIBN, followed by the c-terminal half of the TEV protease^53,54^. The complementary synthetic protein partner contains a Lyn11 membrane tether, CIBN, the optogenetic partner of Cry2PHR, the N terminal portion of the TEV protease, an AsLOV2 domain, which acts to sequester the TEV cleavage sequence while in the dark to reduce any potential dark activity, directly on the N-terminal to a TEV cleavage sequence (ENLYFQ/S), immediately followed by the N1ICD, carrying a C-terminal mEmerald green fluorescent protein tag^56,76^. For the generation of OptoNotch^mut^ a key residue in the canonical TEV cleavage sequence essential for TEV-mediated cleavage was mutated ENLYFQ/S mutated to ENLRFQ/S, which is not susceptible to TEV-induced cleavage^55^.

Sequences for PCR and cloning reactions were acquired from TetO-FUW-N1ICD (addgene,61540), pCMV-NES-CRY2PHR-TevC (addgene,89877), pCMV-TM-CIBN-BLITz1-TetR-VP16 (addgene,89878), PlayBack-CMV-EcoR1 (addgene,203305), Lyn11-GFP-CIBN (addgene,79572), PlayBack-Ef1a-Nde1addgene, 203309, and mEmerald-N1 (addgene,53976).^77^

Hes1-Live-RNA system involves two components: 1) A functional fragment of the human Hes1 promoter, which drives the expression of RNA transcripts carrying 24 copies of the MS2 stem-loop sequence, and 2) a constitutively active cytomegalovirus (CMV) promoter driving the expression of an MS2-coat protein (MCP) ::mScarlet fusion protein ^64,65^.

The Hes1 promoter sequence was made from isolated ShSy-5Y genomic DNA using primers based on the known sequence of the human Hes-1 promoter^64^.

Hes1-Live-RNA (PiggyBac 5” LTR [Hes1-MS2-bGHpa/CMV-mScarlet-MCP-bGHpa/CMV-Puromycin-bGHpa] PiggyBac 3’ LTR) was constructed through a combination of iterative restriction digestions and T4 reactions using MXS cloning as well as Gibson assembly for the final construction of all components into a final plasmid^78^.

EF1α-Live-RNA (PiggyBac 5” LTR [EF1α-MS2-bGHpa/CMV-mScarlet-MCP-bGHpa/CMV-Puromycin-bGHpa] PiggyBac 3’ LTR) was constructed through a combination of iterative restriction digestions and T4 reactions.

The PiggyBac Transposase and pENTR-MCP vectors were gifted by B. Cox (UofT), MS2 24x stem-loop sequences was gifted by Frank Wippich (EMBL), and pmScarlet_C1 was acquired through addgene (85042). All primers used for cloning can be seen in supplementary table 1.

### Cells Lines

HEK293 cells (Cedarlane labs, CRL-1573; RRID:CVCL_0045), HeLa cells (RRID:CVCL_0030) were gifted from Dr. Jeffery Stuart at Brock university, HeLa RBPJ KO cells were gifted from Dr. Tilman Borggrefe (University of Giessen)^16^, and T-ALL [CUTTL1](RRID:CVCL_4966) cells were gifted from Dr. Adolfo Ferrando (Columbia University).

### Cell culture protocol

HEK293 and HeLa cells were cultured in PlasMax media supplemented with 1% pen/strep and 2.5 % fetal bovine serum.^79^ CUTTL1 T-ALL cells were cultured in RPMI-1640 (R0883, Millipore Sigma) supplemented with 2% non-essential amino acids, 20% FBS and 1 % pen/strep. Cells were either cultured on a 35mm collagen coated 1.5 coverslip well plates(P35GCOL-1.5-14-C, Mattek) for live imaging, in-situ hybridization, mRNA isolation and DNA paint experiments; a 24 well uncoated 1.5 coverslip well plate(P24-1.5H-N, Cellvis) for immunohistochemistry, or a 10cm dish for Western blot and protein purification. Cells were grown at 37°C with 5% CO2 in a humidified incubator.

### Cell treatments

#### Transfection

For adherent cells, cells were transfected with Lipofectamine 3000 (Life Technologies, L3000008) following manufacturer’s instruction, and cells were either live imaged or fixed 24 hours post-transfection. For T-ALL cells, cells were transfected using a Neon Transfection System (ThermoFisher) for 3x10ms pulses at 1350 mV. For light-sensitive experiments following transfection, plates were subsequently wrapped in tinfoil and placed in a blackened box inside the incubator.

#### BRDUTP Transfection

Cells immunostained for BRDUTP(Millipore, B0631) were initially treated and transfected with OptoNotch on day 0. The following day, cells were transfected with BRDUTP, and cells were then allowed to incubate for 30 minutes, at which point, cells were fixed and subjected to immunostaining.

#### Drug treatments

Cells treated with either CB103(Selleckchem, S9719) or compound E (ABcam, ab142164) were treated with a 1μM final concentration for 24 hours prior to fixation. If cells were to be transfected and treated, cells were initially treated with compound E or CB103 for 2 hours before transfection, transfected and then either live imaged the following day or fixed the following day. For 1,6-hexanediol (Millipore-Sigma, 240117-50G) treatments, cells were supplemented with 10% of the total volume of the culture media with preheated 50% 1,6-Hexandiol immediately prior to fixation for immunohistochemistry and 3CqPCR or following initial imaging for live cells while still on the microscope.

#### Stable cell production

Hes1-Live RNA stable cells were transfected with PiggyBac [Hes1-MS2-bGHpa/ CMV-mScarlet-MCP-bGHpa/ CMV-Puromycin-bGHpa] along with PiggyBac transposase into HEK293 cells and following 24 hours cells were treated with 1ug/ml Puromycin for 2 weeks changing the media every 2 days. Cells were then taken off of Puromycin for 2 weeks, followed by 2 more weeks of treatment to remove remaining non-stably transfected cells.

#### Nuclear counter staining for live imaging

Cells were treated with 1ul Hoechst 33342(Thermo Fisher, H3570) per 1ml of media for 5min at 37°C prior to live imaging.

#### Live RNA dye imaging

Cells were treated with F22 RNA dye, which was synthesized by Dr John Hayward and John Trant(University of Windsor), at 1μM for 5 minutes and subsequently washed off with fresh media 3 times prior to live cell imaging.^80^

#### OptoNotch Induction

For any OptoNotch experiment not being activated on the microscope itself, cells were illuminated with a white light box for 60 minutes prior to fixation, unless otherwise stated.

### qPCR

RNA was extracted with the Total RNA Purification Kit (Norgen,17200). cDNA was then synthesized from the isolated RNA with iScript™ cDNA Synthesis Kit (Bio Rad) and quantified on a Nanodrop lite instrument (Thermo Fisher). Transcripts were amplified with iQ™ SYBR® Green Supermix (BioRad), and quantitative PCR was performed on a CFX96 real-time qPCR machine (Bio-rad). Primers used can be seen in Table S1. qPCR data was analyzed as fold changes in expression with three separate housekeeping genes as controls. All qPCR experiments included 3 biological replicates that were pooled and measured over 3 technical replicates.

### Antibodies

The following antibodies were purchased from commercial sources : Rat Anti-Notch1(DSHB, BTAN-20; RRID:AB_2153497, 1:50), Rabbit-Anti-Notch(CST, D6F11, 1:200), Rabbit Anti Activated-Notch1(Val1744)(CST, D3B8, 1:1000), Rabbit-Anti-Notch (Atlas,HPA067168; RRID:AB_2685795,1:500), Rabbit Anti-RBPJ (Atlas, HPA060647; RRID:AB_2684337, 1:500), Rabbit Anti-RNAPolII (Atlas, HPA037506; RRID:AB_10672597, 1:500), Mouse Anti-BRD4 (Atlas, AMAb90841; RRID:AB_2665685, 1:500), Rabbit Anti-MED1 (Atlas, HPA052818; RRID:AB_2681962,1:500), Rabbit Anti-P300 (Atlas, HPA004112;RRID:AB_1078746, 1:500), Mouse Anti-BRDUTP (DSHB,G3G4; RRID:AB_1157913,1:1000), Anti-Mouse-568(Invitrogen, A11031, 1:1000), Anti-Rat-568 (Invitrogen, A11077,1:1000), Anti-Rabbit-568 (Invitrogen, A11011, 1:1000), Anti-Mouse-647(Invitrogen, A 21236, 1:1000), Anti-Rat-647 (Invitrogen, A21247,1:1000), Anti-Rabbit-647 (Invitrogen, A21245, 1:1000), Mouse Anti-DIG-568 conjugated(Jackson immuno-research,1:500) .

### SDS-PAGE and Western blot analysis

Unless otherwise stated, samples were lysed in ice-cold RIPA buffer containing 1X protease and phosphatase inhibitors (10 mM phenylmethylsulfonyl fluoride, 1 mM aprotinin, 1 mM sodium orthovanadate and 1 mM sodium fluoride). Samples were homogenized by sonication and briefly centrifuged at 13,000 rpm to remove cellular debris. The concentrations of the resulting protein lysates were determined using the BioRad DC Protein Assay Kit as per manufacturer’s protocol. Unless otherwise stated, all SDS-PAGE was performed on protein lysates using 10% resolving gels and 4% stacking gels run at 80V for 15 minutes and 110V for 90 minutes.

Proteins were then transferred onto 0.2 μm nitrocellulose membranes at 50V overnight (∼16 hours) on ice. Membranes were blocked for 1 hour in a blocking buffer (5% non-fat dry milk in PBS) with constant agitation. Primary antibodies were administered at a dilution of 1:1000 in blocking buffer and incubated overnight at 4°C with constant agitation. Following three washes with PBS + 0.1% Tween 20, membranes were blocked again with blocking buffer for 1 hour and then probed with secondary antibody at a dilution of 1:5000 in blocking buffer for 1 hour at room temperature. To visualize HRP-conjugated secondary antibodies, membranes were probed for 5 minutes with clarity Western enhanced chemiluminescence blotting substrate and imaged with the ChemiDoc Imaging System (Bio-Rad).

### Immunohistochemistry

Cells were fixed in 4% paraformaldehyde in phosphate-buffered saline (PBS) for 10min at room temperature (RT). After three washes in PBS for 5min. Cells were permeabilized with 0.2% triton X100 (Sigma Aldrich, X100) in PBS for 2 min at RT. Following three washes in PBS for 5 min, cells were blocked with either 2% skim milk for antibodies acquired from the DSHB, CST, ABCAM or Invitrogen, or in 4% fetal bovine serum for antibodies acquired from ATLAS antibodies for at least 40 minutes at RT. Primary antibodies were incubated at the previously stated dilution in there given blocking serum for 24 hours at 4°C. Cells were then washed with PBS+0.1% tween (PBST) 3 times. The associated secondary antibody was then incubated on cells at their designated dilution for 2 hours at room temperature in the dark. Cells were washed three times with PBST, and one final 10 min wash in PBS containing 1:1000 Hoechst 33342. Cells were then placed in Vectashield Hardset mounting medium (BioLynx Inc., VECTH1400) and imaged.

### RNA *in-situ* probe synthesis

To produce in-situ probes, cDNA was created identically to our qPCR protocol with the following changes, synthesis was done for 4 hours instead of 1 hour and ethanol precipitation was carried out overnight instead of over the span of 2 hours.. Following cDNA synthesis probe synthesis was carried out as previously described.^81^ With DIG-UTP(SIGMA, DIGUTP-RO), T7 RNA polymerase and RNAPol Reaction Buffer (NEB,M0251) used in our reaction.

### RNA *in-situ* hybridization

Cells were treated with GSI and transfected with OptoNotch. 24 hours post-transfection, cells were fixed and permeabilized as described for immunohistochemical experiments. After fixation cells were placed in hybridization buffer(5% dextran sulfate, 50% formamide, 5× SSC, 100 μg/ml heparin, 100 μg/ml sonicated salmon sperm DNA (Sigma-Aldrich, cat. No. D9156),0.1% Tween-20) for 4 hours at 30 °C. Prior to probe incubation, probes were diluted in hybridization buffer to 1ng/μl, and the solution was incubated at 80°C for 3 minutes, then left on ice for 5 minutes. Probes were then added to cells at 60 °C and hybridized for 24 hours. The following day, cells were washed in 4x SSC for 2 minutes, 2x SSC for 30 minutes, 1x SSC for 30 minutes, and 0.1 x SSC for 20 minutes. Cells were then blocked(2% skim milk,1xPBS, 0.1% tween) for 30 minutes at room temperature. Cells were subsequently stained exactly as described in the immunohistochemistry protocol above.

### Protein purification

Plasmids containing our full-length N1ICD OptoNotch construct were transfected into *HEK293* cells. Following 24 hours post-transfection, they were uncovered and illuminated for 1 hour. Cells were detached using trypsin-EDTA, and N1ICD::GFP was then isolated using GFP-Trap Agarose beads following the prescribed protocol (Chromotek, GTA). Protein concentrations were calculated using a BSA standard curve.

### Droplet assay

Purified proteins were diluted to varying concentrations in buffer containing 50 mM Tris-HCl pHL7.5- and 200-mM glycine with the indicated salt concentrations. 10 μl of each solution was loaded onto an individual uncoated 35mm Dish with a 1.5 coverslip (Mattek, •P35G-1.5-14-C) and imaged.

### Molecular Dynamics Simulations

Molecular dynamics simulations (MDS) were conducted using GROMACS version 2020.1 to simulate the interactions among NOTCH1, RBPJ, MAML1, and a RBPJ DNA binding site.^82,83^ The complete predicted structure files of the NOTCH1 NICD (Valine 1754 to Lysine 2555 at the C-terminus), RBPJ, and MAML1 were obtained from the AlphaFold Protein Structure Database (NOTCH1: AF-P46531-F1-model_v2,; RBPJ: AF-Q06330-F1-model_v2; MAML1: AF-Q92585-F1-model_v2). ^40^ The structure of the human Notch1 transcriptional activation complex), which was derived using only truncated portions of each of Notch1, RBPJ, and MAML1, was extracted from the Protein Data Bank (www.pdb.org; 3v79). We then aligned the complete protein structures from AlphaFold onto the structure of the transcriptional activation complex. Initial MDS resulted in a simulation error in GROMACS due to tight entanglement. Therefore, we manually positioned the proteins in a relatively ‘looser’ position to successfully run the MDS. MDS steps are described as follows; we generated .gro, topol.top, and posre.itp files using a tip3p water model and amber03 force fields. We defined our simulation box using the dodecahedron box type to which we then filled with water, followed by the addition of Na^+^ and Cl^-^ ions to reach a NaCl concentration of 0.15 M and neutralize the system. Following this, we ran energy minimization to ensure that the system had no steric clashes or inappropriate geometry during the MDS. We then performed an equilibration run for 100 ps to bring the system to a temperature fluctuating around 300 K, followed by an equilibration run for 100 ps to bring the system to a pressure of approximately 1 bar. Lastly, the “production” simulation was run for 10 ns (10000 ps). After the MDS was finalized, Visual Molecular Dynamics (VMD) was used to generate the MDS and calculate the root mean square deviation (RMSD) values relative to the starting frame.^84^

### 3C PCR/qPCR

3C-PCR was completed as previously described using primers previously designed to study the interaction between the MYC and the NDME locus (Table S1).^8,50–52,68^ Analysis was completed using qPCR and agarose gel images to show representative band produced after qPCR reaction. Analysis was completed whereby the signal produced by MYC-NDME was divided by the signal of MYC-MYC to get a relative association rate. This was then done for the negative control MYC-NDME+80K from the same sample to act as a random association control.

### DNA-PAINT

DNA PAINT was completed as previously described with the following modifications.^70^ Cells were instead grown on a 35 mm matek collagen-coated plate and incubated in excess volume of staining solution to remove the requirement for sealing with rubber cement. Once cells were stained, they we mounted in Vectashield Hardset mounting medium and imaged. Probes were designed to target either the MYC locus(chr8: 127730000-127740000) or the NDME locus(chr8: 130175000-130185000). MYC Probe barcodes were conjugated to cy3 and NDME probe barcodes were conjugated to Cy5(IdtDNA) (Table S1).

### Microscopy

All imaging was conducted on an inverted Zeiss Axio Observer spinning disc confocal microscope equipped with a Yokogawa spinning disc head and a Prime BSI 16-bit camera fitted with 4 laser lines (350-400 nm, 450-490 nm, 545-575 nm, 625-655nm) and a Zeiss Direct FRAP FLIP Laser Manipulation for Axio Observers (Zeiss,423635). Most imaging was completed with a 40x 1.4 NA Plan Apochromat oil objective, except for droplet assay imaging which used a 20x 0.8 NA Plan Apochromat air objective. Imaging was conducted on a stage-fitted dark box incubator with CO2 and temperature regulation to allow live imaging at 37°C with 5% CO2. Image analysis was subsequently carried out on ImageJ(RRID:SCR_002798) as described below.^85^

### Image analysis

#### Condensate volume and counting analysis

Nuclei of individual cells were isolated, using Hoechst as an ROI to isolate signal only emitting from the nuclei of individual cells. We then applied the 3d object counter function in ImageJ to determine the number and volume of each condensate within a given nucleus.

#### Nuclear localization rate

We first isolated nuclear proteins as described above in *Condensates volume and counting*. The change in Nuclear GFP signal was measured at each time point and then normalized to time 0 within each condition, and these values were then averaged across multiple trials to determine rate of N1ICD nuclear translocation following OptoNotch activation.

#### Colocalization analysis

We first Isolated for nuclear proteins, as done above in *condensates volume and counting*, we then calculated Manders coefficients for N1ICD signal and the given co-stain to determine total nuclear localization between the two in relation to total nuclear N1ICD signal. We then measured the total amount of fluorescence of the co-staining protein within the nucleus and then measured the amount of signal that colocalizes with nuclear N1ICD and represented that as a fraction of total nuclear protein. To then get the ratio of N1ICD condensates that contained some level of a co-staining protein we then took a total count of the number of nuclear N1ICD condensates and compared that to the number of N1ICD condensates that contained some level of co-staining protein.

#### FRAP analysis

Using the Zeiss Direct FRAP FLIP Laser Manipulation for Axio Observers, one of two different conformations of photo-bleaching areas were used to either: bleach a single Nuclear N1ICD condensates, or bleach a partial area of a single condensates along with the surrounding area. Bleached areas were measured every 2 seconds following bleaching or imaged every 30 seconds for SRRF data collection. Bleach Frap kinetics were fit using R to the first order kinetic in quantifying our FRAP data, we fit the average bleach kinetic to the equation.

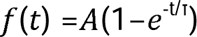

where A represents the mobile fraction, τ is the time constant, and t is the time post photo-bleaching.

#### SRRF image acquisition

To acquire SRRF data the capture field of the camera was reduced to 400x400 pixels and 200 images from a single optical field of view were acquired at 100 % laser power with an excitation time of 500 μs. SRRF images were then calculated using the NANOJ SRRF plugin in ImageJ.^62,63^ To validate our SRRF outputs, a random subset of 250 images across all conditions were chosen to be run through squirrel analysis for accuracy.^61^ In order to collect Z-Stacks of SRRF images we performed SRRF imaging on a single optical plane followed by a single step of 200nm to the next optical plane to which we would then perform SRRF imaging, this was repeated through the stack up to 12 SRRF images. All SRRF images shown in figures are of single optical sections. To produce 3D renders for videos 3D viewer in ImageJ was used.

#### SRRF arrangement determination

After identifying individual nuclear N1ICD condensates ROIs were drawn around all unique fluorescence signal of the co-staining protein within a condensate. The minimal distance between the centre of mass of each ROI and the nearest edge of the encapsulating N1ICD shell was measured.

#### FRAP Distance Recovery

*N*1ICD condensates that were half photo-bleached *and imaged with* SRRF were skeletonized and the max skeleton length of each image was measured and the change in length was calculated to determine a nm/sec travel rate for proteins within a given condensates.

#### FRAP Percent Area Recovery

Individual Pre-Bleach Condensate SRRF images were measured for total fluorescence and normalized to 1, and total fluorescence for all subsequent SRRF images was measured and represented as a ratio of the initial intensity as a metric of the total area recovered in individual condensates. This was then repeated for each time point and averaged across trials.

#### Analysis of N1ICD Condensate Fusion

ImageJ TRACKmate was used for detecting the movement of condensates within a cell.^86^ To isolate for fusion events condensates that intercepted with one another, leaving only an individual condensate, were isolated and subsequently quantified prior to and following fusion to determine the volume and total fluorescence over time.

##### MS2-Live-RNA Distance Analysis

The centre of mass for each MS2-Live-RNA focus and N1ICD condensate was recorded for each a field of view. We then analyzed each MS2-live RNA’s centre of mass and determined the distance to the nearest NICD condensate.

#### MS2 bursting Analysis

For this analysis, we created three separate categories of ROIs to bin OptoNotch condensates and Hes1-Live-RNA transcriptional foci into, these being: N1ICD condensates that had mScarlet signal above a detection threshold, NICD foci that did not have mScarlet signal above a detection threshold and Hes1-Live-RNA signal that was produced in the absence of OptoNotch. The ROIs were identified and tracked using ImageJ, and the maximal fluorescence intensity values for both channels within each ROI at each time point were recorded. To interpret the relationship between NICD and Hes1-Live RNA for each ROI, we produced Z-score for each channel and heat-mapped those points with respect to time post OptoNotch activation. This was done to see a correlation between the formation of NICD condensates and the occurrence of Hes1-Live-RNA foci. By then recording the duration that Hes1-Live-RNA signal stayed above the threshold for any given condensates, we were able to determine the duration of signalling either with or without the co-occurrence of NICD.

#### Nearest Neighbor Calculation for DNApaint

MYC and NDME nuclear foci were thresholded, and each position was labelled as the centre of fluorescence for each focus. We then isolated each nucleus in the field of view and measured the minimal distance between the MYC and NDME loci within each nucleus. For the determination of the role of OptoNotch, prior to distance calculation, OptoNotch signal was used as an ROI only to include signal colocalizing with OptoNotch condensates, and the distance between the MYC and NDME loci was subsequently quantified across conditions.

#### Statistics and Data Analysis

All experiments were completed with a minimum of 3 separate technical replicates comprising 3 biological replicates (a minimum of nine total plates/wells with three plates imaged at a time repeated over three separate experiments). The individual number of measurements (n’s) of each experiment is represented in each corresponding figure caption. All statistical analyses were completed using R studio.^86^

#### H-IC data analysis

Hi-C heatmap was generated using Juicebox based on user manual at 10Kb resolution. T-ALL data used was taken from Juicebox’s publicly available repository of Hi-C sequenced cell lines.^87^

## Acknowledgments

This study was supported by the Natural Sciences and Engineering Research Council of Canada Discovery Grants(2018-06781) and NIH grant R35GM133482. We also acknowledge Dr’s Brian Cox(UofT), Frank Wippich(EMBL), Jeffery Stuart(Brock University), Rebecca MacPherson(Brock University), Tilman Borggrefe (University of Giessen), Adolpho Ferrando (NYU Langone), Yi Sheng(York University), and John Trant and John Howard(University of Windsor) for various reagents and help provided throughout the development of this manuscript.

## Author contributions

Conceptualization, G.F and A.N.; Methodology, G.F and A.N.; Investigation, G.F., R.D.H., M.M., and A.T.; Writing – Original Draft, G.F., and A.N.; Writing – Review & Editing, G.F., R.D.H., M.M., A.T., D.A., Y.L., V.L., and A.N.; Funding Acquisition, A.N. and V.L. ; Formal Analysis, G.F. and Y.L.; Resources, D.A., V.L., and A.N.; Supervision, A.N.

## Data Availability

All data presented in this study are available within the manuscript and supplementary material.

## Materials & Correspondence

For all correspondence, please contact Aleksandar Necakov

---

**Extended Data Figure 1: Molecular Dynamic Simulation Of The Notch1 Intracellular Domain (AA1754-AA2555)**

A) Representative heatmap of the root means squared deviation(RMSD), in angstroms(Å), of individual residues following molecular dynamic simulation in GROMACS2 with RMSD calculated in VMD over 2000 frames spread over the simulation. B) Domain RMSD plots over time looking at the structural movement of either the RAM and Ankyrin repeat domain(1754-2119, blue), the TAD domain and c-terminal tail(2120-2555, green), or the entirety of the Notch1 intracellular domain(1754-2555, purple) C) N1ICD structural analysis using Prediction of prion-like domains (PLD), Net charge per residue (NCPR), Fraction of charged residues (FCR), and hydrophobicity analysis.

**Extended Data Figure 2: Effect of 1,6-Hexanediol treatment on Notch1 Condensates**

A) Formation of hollow cavities(arrows) in *in-vitro* droplets at 250mM NaCl. Brightfield(top), green fluorescence (bottom). Scale Bar 10μm. B) Isolated NICD::GFP treated either with control (left) or 1,6-Hexanediol treatment, concentration used was 100um NICD::GFP in 150mM NaCl. Scale Bar 50μm C) Live imaging of N1ICD in OptoNotch activated HEK293 cells before and six seconds after treatment with either vehicle(DMSO) or 1,6-Hexandiol. Scale bar 10μm. D) Quantification of the number of N1ICD foci per nucleus in cells from panel C, showing that 1,6-Hexandiol treatment results in a near-complete loss of intranuclear N1ICD foci from 29.7(+/- 30.1) in vehicle control cells to 0.6(+/-0.8) in 1,6-Hexanediol-treated cells. N=1000 cells per condition. E) Quantification of the total intensity of Notch1 in OptoNotch-activated HEK293 cells from panel C, showing that treatment with 1,6-Hexanediol does not significantly impact total Notch1 intensity in comparison to vehicle control treated cells, 2643(+/-877) and 2566(+/-599) A.U of Notch1, respectively. N= 1000 cells per condition. $<0.1 student t-test. See Source Data Extended Figure 2.

**Extended Data Figure 3: Antibody validation and comparison of commercial Notch1 antibodies**

A) Notch1 immunostainings from HEK293 cells either plated on control(top), or DeltaMAX(bottom) using CST antibody. B) Total nuclear Notch1 protein from panel A showing control and DeltaMAX treated cells respectively had 946(+/-312) and 1276(+/-462) A.U of Notch1. N= 5000 nuclei per condition. C) Volume of individual nuclear foci from panel A showing control and DeltaMAX treated cells respectively form Notch1 foci that are on average 0.141(+/-0.153), and 0.554(+/-0.335). N= 51000 foci per condition. D) Total Individual Fluorescence per foci from panel A showing control and DeltaMAX treated cells respectively have foci that have 37(+/-38.7) and 65.6(+/-40.1) A.U of Notch1. N= 51000 foci per condition. E) Notch1 immunostainings from HEK293 cells either plated on control (Top) or DeltaMAX (Bottom) using DSHB antibody. F) Total Nuclear Notch1 protein per Cell from panel E showing control and DeltaMAX treated cells respectively had 1891.5(+/-581.1) and 3194.3(+/-3343.9) A.U of Notch1. N= 3500 nuclei per condition. G) Volume of individual nuclear foci from panel E showing control and DeltaMAX treated cells respectively form Notch1 foci that are 0.142 μm^3^ (+/-0.238) and 0.235 μm^3^ (+/-0.635) average. N= 4000 foci per condition. H) Total Individual Fluorescence per foci from panel E showing control and DeltaMAX treated cells respectively have foci that have 69.1(+/-109.7) and 99.3(+/-249.88) A.U of Notch1. N= 4000 foci per condition. I) Endogenous Notch1 fluorescence immunostainings in T-ALL cells treated either with DMSO or GSI using CST antibody. J) Total amount of Nuclear Notch1 protein per cell from panel I showing DMSO and GSI treated cells have 1159(+/-507) and 774(+/-460) A.U of Notch1, respectively. N= 31000 cells measured per condition. K) Volume of individual nuclear foci from panel I showing DMSO and GSI treated cells form Notch1 foci that are on average 0.658μm^3^(+/- 0.636) and 0.236μm^3^(+/-0.127) respectively. N= 54000 foci measured per condition. L) Total Fluorescence per individual nuclear foci from panel I showing DMSO and GSI treated cells have foci of 57.9(+/-65.3) and 25.6 (+/-14.2) A.U of fluorescence intensity for Notch1. N= 54000 foci measured per condition. M) Endogenous Notch1 fluorescence immunostainings in T-ALL cells treated either with DMSO or GSI using DSHB antibody. N) Total amount of Nuclear Notch1 protein per cell from panel M showing DMSO and GSI treated cells have 3516(+/-1571) and 1556(+/-680) A.U of Notch1, respectively. N= 4000 cells measured per condition. O) Volume of individual nuclear foci from panel M showing DMSO and GSI treated cells form Notch1 foci that are on average 0.564μm^3^(+/-0.502) and 0.235μm^3^(+/-0.128) respectively. N= 12000 foci measured per condition. P) Total Fluorescence per individual nuclear foci from panel M showing DMSO and GSI treated cells have foci of 40.5(+/-43.7) and 21.1(+/-11.6) A.U of fluorescence intensity for Notch1. N= 12000 foci measured per condition. *p<0.01 One-way ANOVA+Tukey post-hoc, $p<0.01 student t-test. See Source Data Extended Figure 3

**Extended Data Figure 4: Cleavage of Notch1 in the absence and presence of exogenous, surface-immobilized DeltaMAX ligand:**

A) Western blot of HEK293 cells following nuclear/cytoplasmic fractionation in the absence or presence of GSI inhibitor. Activated Notch1 antibody staining shows cleavage of endogenous Notch1(left two lanes), with cleaved Notch1 present at high levels in the nucleus compared to the cytoplasm. GSI treatment inhibits Notch1 cleavage, as shown by the loss of the activated Notch1 band(Right two lanes). Lamin A/C and β–tubulin was used as loading controls for the nuclear and cytosolic fractions, respectively. B) Western blot of HEK293 cells cultured either in the absence of exogenous ligand (control) or with immobilized DeltaMAX ligand. Activated Notch1 antibody staining shows a band corresponding to cleaved endogenous Notch1, which is present at high levels in the nucleus compared to the cytoplasm (control, lanes one and three). Treatment with surface-immobilized DeltaMAX ligand activates Notch1 cleavage over baseline control, as shown by the increase in the abundance of activated Notch1 in the nuclear fraction(DeltaMAX, lanes two and four). Lamin A/C and β–tubulin was used as loading controls for the nuclear and cytosolic fractions, respectively. N=3 for each blot.

**Extended Data Figure 5: Comparison of OptoNotch condensate properties in untreated and DeltaMAX-activated cells.**

A) Immunostaining against Notch1 in T-ALL cells, DeltaMAX-plated HEK293 cells, and GSI-treated OptoNotch activated HEK293 cells using a secondary 647 that has been re-colourized to green. Scale bar 10μm. B) Quantification of Hes1 expression by qPCR in either GSI treated, DeltaMAX plated, or OptoNotch activated HEK293 cells, showing a -0.736(+/-0.839), 13.777(+/- 0.850), and 6.465(+/-1.385) fold change in expression, respectively. C) Quantification of Total Nuclear Notch1 showing T-ALL cells, DeltaMAX plated HEK293 cells, or OptoNotch activated HEK293 cells have 1782(+/-2137),1463(+/-1005) and 3841(+/-3181) A.U of Notch1, respectively. N=3000 cells per condition D) Quantification of the total volume per foci from T-ALL cells, DeltaMAX plated HEK293 cells, or OptoNotch activated HEK293 cells form Notch1 foci that are on average 0.640 μm^3^ (+/-0.506), 0.551 μm^3^ (+/-0.343), and 0.554 μm^3^ (+/-0.406), respectively. N= 5000 foci per condition. E) Quantification of the total Intensity of Individual Notch1 foci from T-ALL cells, DeltaMAX plated HEK293 cells, or OptoNotch activated HEK293 cells form Notch1 foci that have an average fluorescence intensity of 26.1(+/-49.8), 18.3(+/- 37.1), 23.3(+/-56.4) A.U., respectively. N= 5000 foci per condition. F) Quantification of the total number of Notch1 foci per cell showing T-ALL cells, DeltaMAX plated HEK293 cells, or OptoNotch activated HEK293 cells form 8.23(+/-4.6), 17(+/-21.4), and 32.2(+/-29.3) Notch1 Foci per cell, respectively. N= 3000 cells per condition. *p<0.01 One-way ANOVA+Tukey post-hoc. See Source Data Extended Figure 5.

**Extended Data Figure 6: Super-Resolution Quantitative Image Rating And Reporting Of Error Locations(SQUIRREL) Analysis**

A) Original representative confocal image acquired using spinning disc confocal microscopy (Top left). Post-SRRF analysis of confocal images (Top right). Convolved SRRF image for error mapping of confocal image (Bottom left). Error map of SRRF data showing a Resolution Scaled Pearson’s(RSP) of 0.974 and an average resolution scaled error (RSE) of 3.732 (Bottom right). Scale bar 5 μm. B) Dot plot of average RSE value output by individual SRRF images. N=250. C) Dot plot of RSP value output by individual SRRF images. N=250 See Source Data Extended Figure 6.

**Extended Data Figure 7: Additional T-ALL N1ICD Molecular Condensates SRRF Imaging .**

T-ALL cells immunostained against Notch1 Imaged using SRRF microscopy showing the formation of ring-like structed N1ICD condensates. Scale Bar 500nm

**Extended Data Figure 8: Addition SRRF Analysis of Notch1 Nuclear Condensates**

A) Direct size comparison between an initial focus (pre-bleaching, pink) to its final state following photo-bleaching recovery from Figure 3.F. B) Direct size comparison between an initial focus (pre-bleaching, pink) to its final state following photo-bleaching recovery from Figure 3.G. C) SRRF images of an intranuclear N1ICD focus showing a decrease in size over time. D) SRRF images of an intranuclear N1ICD focus showing an increase in size over time. E) Overlay of the initial (0 minutes; pink), and final image (7 minutes; green) from Panel C. F) Overlay of the initial (0 minutes; pink), and final image (7 minutes; green) from Panel D. G) Growth evolution of an N1ICD condensate using SRRF imaging, clearly demonstrating the progressive formation of a hollow core over time (180 minutes). H) Boxplot of Endogenous and OptoNotch Notch1 condensate signal widths with average widths of 194nm (+/-112) and 198nm (+/-66) respectively. N=100 condensates per condition. Scale bar 500nm. See Source Data Extended Figure 8

**Extended Data Figure 9: Non-Transfected Control cells from Figure 4 panel A:**

Immunofluorescence images of HEK293 treated with GSI staining for either RBPJ, MAML1, p300, MED1, BRD4, RNAPOLII, or BRD-UTP. Scale Bar 10 μm.

**Extended Data Figure 10: RNA Polymerase 2 colocalizes with OptoNotch Condensates In Living Cells.**

A) HEK293 cells were transfected with either DENDRA-RNAPolII (top panel), activated OptoNotch (middle), or both activated OptoNotch and DENDRA-RNAPolII (bottom) and imaged prior to UV photoconversion. B) Cells from Panel A post UV photoconversion with both 488 and 568 nm light, showing strong colocalization between green and red signals in intranuclear N1ICD condensates. C) Quantification of the ratio of red to green fluorescence in each condition prior to UV photoconversion N=60 cells per condition. D) Quantification of the ratio of red to green fluorescence in each condition following UV photoconversion N=60 cells each per condition. E) Live fluorescence image of a GSI treated, OptoNotch (green) activated, HEK293 cell stained with live RNA dye (purple). Scale bar 1 μm. A/C Scale bar 10μm, E Scale bar 2 μm. *p<0.01 One-way ANOVA with Tukey post hoc. See Source Data Extended Figure 10.

**Extended Data Figure 11: OptoNotch activates Notch target genes Hes5 And Hey1**

A) Quantification of Hes5 expression by qPCR in wildtype, OptoNotch activated, GSI-treated, or OptoNotch activated and GSI-treated cells, showing a relative expression of 1(+/- 0.440),14.913(+/-4.265), 0.0947(+/-0.0765), and 7.673(+/-0.639) for Hes5 respectively. B) Quantification of Hey1 expression by qPCR in wildtype, OptoNotch activated, GSI-treated, or OptoNotch activated and GSI-treated cells, showing a relative expression of 1(+/-0.651), 10.139(+/-2.887),0.00341(+/-0.0176), and 7.205(+/-1.173) for Hey1, respectively. Bar Graph represents mean +/- SD. All data acquired in HEK293 cells. *p<0.01 One-way ANOVA with Tukey post hoc. See Source Data Extended Figure 11

**Extended Data Figure 12: Replicates of SRRF Co-Localization from Figure 4 I.**

A) RBPJ containing condensates. B) MAML1 containing condensates C)p300 containing condensates D) BRD4 containing condensates E) RNA PolII containing condensates F) MED1 containing condensates G) BRD-UTP containing condensates. Scale Bar 500nm.

**Extended Data Figure 13: Validation of Stably transfected Hes1-Live-RNA MS2.**

A) HEK293 cells stably transfected with the Hes1-Live-RNA reporter, treated with either vehicle (DMSO), GSI, or GSI in conjunction with OptoNotch activation. Scale bar 5 μm. B) Quantification of the number of Hes1-Live-RNA foci per cell in cells treated with either vehicle control, GSI, or, GSI in conjunction with OptoNotch activation, showing 17.4(+/-8.1), 5.1(+/-3.6), and 20.1(+/-12) Hes1-Live-RNA foci per cell, respectively. N=300 cells per condition. C) Example of endogenously activated Stably transfected Hes1-Live-RNA cells. Scale bar 5 μm. See Source Data Extended Figure 13.

**Extended Data Figure 14: ΔANK-N1ICD condensates similarly form hollow condensates.**

A) Matched confocal and SRRF microscopy of ΔANK-N1ICD transfected HEK293 cell. Scale bar 10μm B) Insets of three ΔANK-N1ICD condensates showing hollow core formation under SRRF microscopy. Scale bar 500nm.

**Extended Data Figure 15: N1ICD condensates contain RBPJ and MAML1: two core components of the Notch transcriptional activation complex.**

A) Fluorescence immunohistochemical staining of HEK293 cells expressing ΔANK-N1ICD, showing prominent condensates co-stained for either RBPJ (top) or MAML1(bottom). Scale bar 10μm B) Zoom insets from Panel A showing individual condensates. Scale bar 2μm C) Quantification of the proportional number of ΔANK-N1ICD condensates that contain any amount of either RBPJ 0.816(+/-0.156), or MAML1 0.933(+/-0.0523). N= 250 cells per condition. $ p<0.01 on Student T-test. See Source Data Extended Figure 15

**Extended Data Figure 16: MYC RNA expression in HEK293 cells.**

A) RNA *in* situ hybridization of MYC RNA in combination with fluorescence immunostaining against Notch1 protein in either DMSO control, GSI-treated, or OptoNotch activated cells. Scale bar 10μm.B) Comparison of the total Nuclear MYC RNA fluorescence in either DMSO, GSI or OptoNotch activated cells showing and average of 3640(+/-2878), 198(+/-329), and 4235(+/- 4231) AU of MYC RNA, respectively. N = 600 cells per condition. C) Quantification of MYC expression by qPCR in HEK293 cells treated with either vehicle, GSI or GSI+OptoNotch, showing a relative expression of MYC of 1.00(+/-0.136),0.0835(+/-0.0380), and 2.889(+/-0.881), respectively for each condition. *p<0.01 One-way ANOVA with Tukey post hoc. See Source Data Extended Figure 16

Supplemental table 1: Primers and oligonucleotides used.

Movie 1:OptoNotch Activation Leading To Subsequent Nuclear Localization Of N1ICD. In HEK293 Cells Imaged Over 25 Minutes. Scale Bar 10um.

Movie 2 : N1ICD Condensate Photo-bleaching Video Over 5 Minutes. Photo-bleached area Is Encircled In Yellow.

Movie 3 : Two N1ICD Condensates Undergoing Fusion Over The Time Course Of 5 Minutes. Scale bar 1 micron

Movie 4: 3D structure of Endogenous Notch1 condensates in T-ALL cell. Render facing the X-Y plane while rotating around the Y axis(left), Render Facing the X-Z plane while rotating around the Z axis(centre), SRRF images used to produce the render(right). Scale in left and centre is in um, scale bar 500nm.

Movie 5: 3D structure of OptoNotch condensates in HEK293 cell. Render facing the X-Y plane while rotating around the Y axis(left), Render Facing the X-Z plane while rotating around the Z axis(centre), SRRF images used to produce the render(right). Scale in left and centre is in um, scale bar 1 um.

Movie 6 : N1ICD Condensate Undergoing Growth Phase Showing An Increase In The Total Volume Of A Single Condensate Over 16 Minutes. Volume On Condensate Indicated In Centre Of Condensate Measured For Each Frame. Scale bar 1 micron

Movie 7 : ΔTAD-N1ICD Condensate photo-bleaching Video Over 5 Minutes. Bleach Area Is Encircled In Yellow.

Movie 8 : ΔAnkyrin-N1ICD Condensate photo-bleaching Video Over 5 Minutes. Bleach Area Is Encircled In Yellow.

